# Fungal-derived tRNAs are expressed and aminoacylated in orchid mitochondria

**DOI:** 10.1101/2024.11.19.624230

**Authors:** Jessica M. Warren, Luis F. Ceriotti, M. Virginia Sanchez-Puerta, Daniel B. Sloan

## Abstract

Plant mitochondrial genomes (mitogenomes) experience remarkable levels of horizontal gene transfer (HGT), including the recent discovery that orchids anciently acquired DNA from fungal mitogenomes. Thus far, however, there is no evidence that any of the genes from this interkingdom HGT are functional in orchid mitogenomes. Here, we applied a specialized sequencing approach to the orchid *Corallorhiza maculata* and found that fungal-derived tRNA genes are transcribed, post-transcriptionally modified, and aminoacylated. In contrast, all the transferred protein-coding sequences appear to be pseudogenes. These findings show that fungal HGT has altered the composition of the orchid mitochondrial tRNA pool and suggest that these foreign tRNAs function in translation. The exceptional capacity of tRNAs for HGT and functional replacement is further illustrated by the diversity of tRNA genes in the *C. maculata* mitogenome, which also include genes of plastid and bacterial origin in addition to their native mitochondrial counterparts.

## Introduction

Plant mitochondrial genomes (mitogenomes) are evolutionary hotspots for horizontal gene transfer (HGT), with high rates of organelle fusion and recombination resulting in transferred sequences from diverse sources, including plastids, the mitochondria of other plants, and even bacteria (Rice et al. 2013; Knie et al. 2015; Sanchez-Puerta et al. 2017). Although most transferred sequences are likely non-functional and readily lost, tRNA genes are exceptional in that they have frequently gained function in the recipient mitogenome (Joyce and Gray 1989; Marchfelder et al. 1990; Kitazaki et al. 2011), sometimes replacing the native mitochondrial copy (Marchfelder et al. 1990; Duchêne and Maréchal-Drouard 2001; Warren and Sloan 2020). Recently, another source of HGT was discovered with the report that fungal mitochondrial DNA has been inserted into the mitogenome of orchids (Sinn and Barrett 2020). A small insertion of ∼270 bp occurred prior to the divergence of the orchid family 76-84 Mya and was later replaced by a larger (∼8 kb) insertion in the epidendroid orchids 28-43 Mya (Ramírez et al. 2007; Valencia-D et al. 2023). Together, these fungal-derived sequences contain seven mitochondrial tRNA (mt-tRNA) genes. Although expression of these genes was not detected (Sinn and Barrett 2020), tRNAs are notoriously difficult to sequence with conventional RNA-seq methods (Wilusz 2015; Padhiar et al. 2024). Therefore, it remains unclear whether this case of interkingdom HGT has functionally altered the make-up of the orchid mitochondrial tRNA pool. To address this question, we applied a specialized tRNA-seq strategy (Watkins et al. 2022) that enables robust detection of tRNAs to the coralroot orchid *Corallorhiza maculata* (Figure 1a), a non-photosynthetic species that relies entirely on mycorrhizal fungi for carbon acquisition (mycoheterotrophy). We coupled this approach with a chemical pretreatment (sodium periodate followed by sodium tetraborate) that distinguishes between aminoacylated and uncharged tRNAs based on whether an amino acid is present to protect the tRNA from removal of its terminal 3′ nucleotide (Evans et al. 2017; Davidsen and Sullivan 2024). Determining the aminoacylation state for the foreign mt-tRNAs is essential for understanding whether they are recognized and charged by orchid aminoacyl-tRNA synthetases (aaRS) and, thus, have the potential to function in translation.

**Figure 1.**
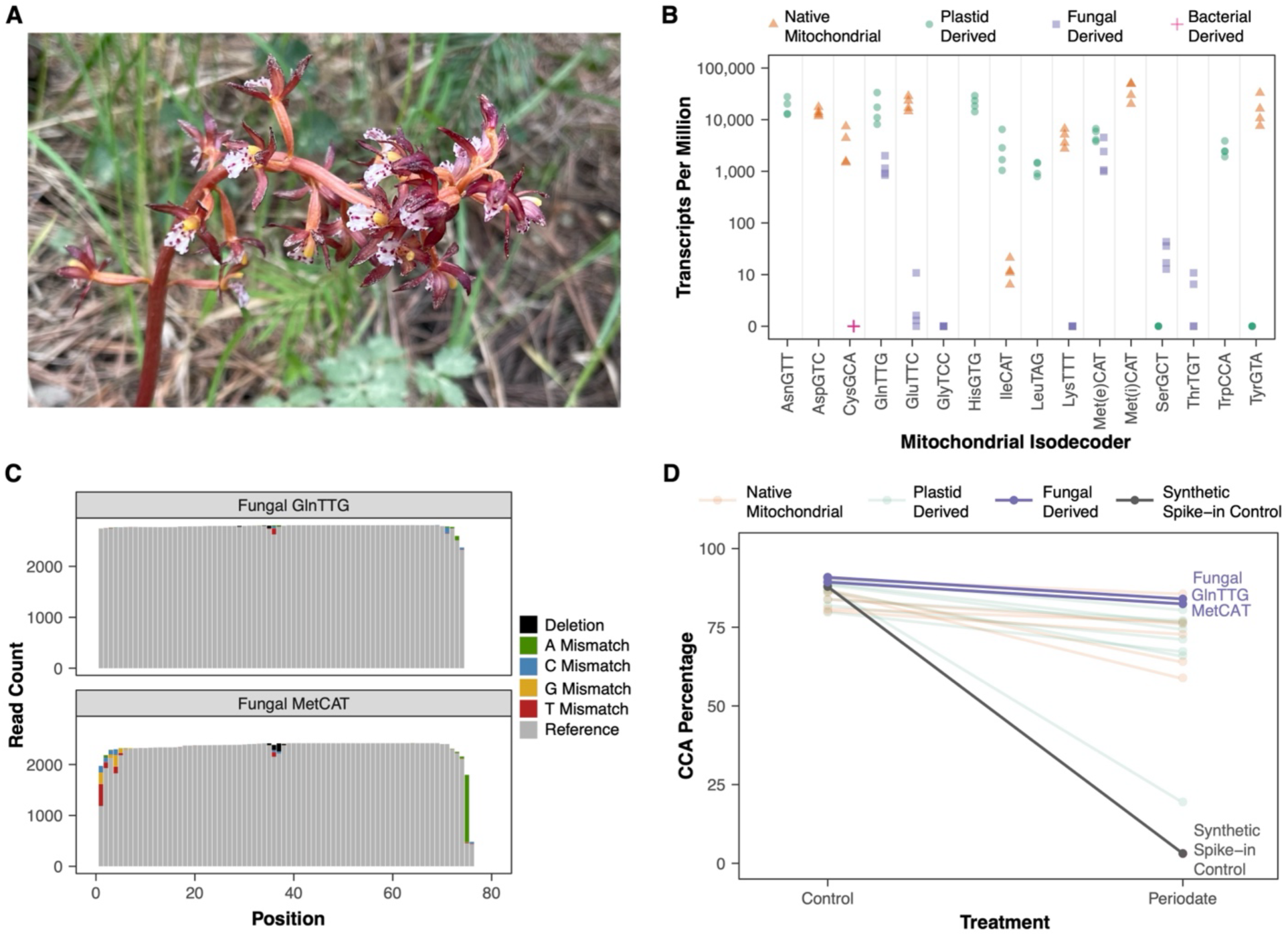
Expression, post-transcriptional modification, and aminoacylation of some fungal-derived tRNA genes in the *Corallorhiza maculata* mitogenome. (A) Photograph of one of the *C. maculata* individuals used in this study. (B) Expression of mitochondrial-encoded tRNAs. Read abundances for each of four *Corallorhiza maculata* biological replicates (two flowers from each of two plants) are quantified as transcripts (reads) per million mapped to organellar reference sequences. The origin of mitochondrial tRNA genes is indicated by point shape and color. Data are only plotted from control (no periodate) libraries and for reads that mapped uniquely to a reference sequence. Values for three different plastid-derived tRNA-IleCAT genes are summed (Table S1). Two of these are identical copies that account for 85% of the reads. The third differs by only a single substitution at the discriminator base position and may be functioning as a t-element (Forner et al. 2007) involved in 3′ processing of *nad7* rather than in translation (see Figure S2; Scaffold 3). Met(e)CAT and Met(i)CAT refer the elongator and initiator tRNA-Met genes, respectively. The fungal-derived MetCAT was arbitrarily included in the Met(e)CAT category, as the initiator/elongator distinction does not apply to fungal mitogenomes. Four plastid-derived tRNA genes in the *C. maculata* mitogenome (AsnGTT, CysGCA, PheGAA, and ThrTGT) were excluded from the reference because they are identical in sequence to their plastid-encoded counterparts and, therefore, indistinguishable in mapping. (C) Summary of read coverage and sequence variants in the two fungal-derived tRNA genes with substantial read abundance (GlnTTG and MetCAT). Raw read counts are summed across the four control (no periodate) libraries after excluding reads that are truncated at the 3′ end (lacking > 7 nt). Deletions and nucleotide mismatches are indicated by color. The lack of major internal drops in coverage indicates that RT was not inhibited by “hard-stop” base modifications in these two tRNAs. The cluster of deletions and misincorporations at position 37 in both tRNAs suggests base modifications immediately 3′ of the anticodon. The high frequency of A mismatches at the second-to-last nucleotide position in tRNA-MetCAT reflects the fact that many of its transcripts are modified with a CA tail rather than a full CCA tail. The abundance of mismatches at the 5′ end of tRNA-MetCAT may reflect artefactual addition of nucleotides by the reverse transcriptase after reaching the end of transcripts that are shorter than the annotated reference sequence. (D) Periodate treatment indicates high levels of aminoacylation for most expressed mitochondrial tRNAs, including the fungal-derived GlnTTG and MetCAT genes. Reported values represent the percentages of reads with intact CCA tails after excluding reads that lacked more than just a single 3′ nucleotide. Values were averaged across four biological replicates. Only genes with >30 reads per library are shown. The synthetic spike-in control tRNA shows that tRNAs that are not protected by an amino acid experience near complete removal of 3′ nucleotides in response to periodate. The retention of intact CCA tails following periodate treatment in most of the mitochondrial tRNAs (including the two fungal-derived tRNAs with substantial expression levels) indicates high levels of aminoacylation. The one notable exception with little retention of the 3′ nucleotide is the plastid-derived tRNA-IleCAT that may be acting as a *nad7* t-element (see above). In contrast, transcripts associated with the other plastid-derived tRNA-IleCAT retain intact CCA tails at levels comparable to other mt-tRNAs. The fungal-derived tRNA-Met percentages do not include reads that were only post-transcriptionally modified with a CA addition (instead of CCA). The CA-tailed reads exhibited much lower retention of the 3′ nucleotide after periodate treatment (Table S2).

### The Corallorhiza maculata mitogenome contains tRNA genes of diverse origins

The assembled mitogenome of *C. maculata* is a posterchild for tRNA HGT, with 16 plastid-derived, one bacterial-derived, seven fungal-derived, and seven native mt-tRNA genes (Table S1). The one bacterial gene (tRNA-Cys) has been identified in numerous angiosperms, including some in which it is expressed and aminoacylated (Kitazaki et al. 2011). However, we did not detect any expression of this gene in *C. maculata* (Figure 1b). The plastid-derived mt-tRNA gene set represents 12 different anticodons and includes many genes that were anciently transferred during seed plant evolution (Richardson et al. 2013). It also includes expressed tRNAs that are not typically plastid-derived (or in some cases present at all) in plant mitogenomes, including tRNA-Gln, tRNA-Ile, and a copy of tRNA-Leu with partially disrupted base-pairing in its acceptor stem. We detected expression of all seven of the native mt-tRNAs. Although raw read abundances from tRNA-seq data should be interpreted cautiously due to large biases in the sequencing process (Warren, Salinas-Giegé, Hummel, et al. 2021), it is noteworthy that the native tRNA-Ile was only detected at very low levels, whereas transcripts from the plastid-derived tRNA-Ile genes were abundant (Figure 1b). This observation suggests that *C. maculata* may be undergoing a functional replacement of the native tRNA-Ile with its plastid-derived counterpart. Despite the gain of so many mt-tRNA genes via HGT, *C. maculata* does not contain a minimally sufficient set of tRNA genes in its mitogenome to support translation and must presumably import nuclear-encoded tRNAs from the cytosol like other angiosperms (Schneider 2011). In contrast to the complex mix of mt-tRNA genes, the *C. maculata* plastome has the conventional angiosperm set of 30 tRNAs, all of which appear to be expressed at substantial levels (Figure S1). Therefore, despite being an obligate heterotroph (i.e., non-photosynthetic), *C. maculata* shows no signs of the plastid tRNA gene loss that is often observed in plants with a more ancient history of heterotrophy (Wicke and Naumann 2018).

### Fungal-derived tRNA genes in the Corallorhiza maculata mitogenome are transcribed, post-transcriptionally modified, and aminoacylated

Our tRNA-seq analysis detected transcripts from five of the seven fungal-derived tRNA genes, although only two of them (tRNA-Gln and tRNA-Met) had substantial read abundances (Figure 1b). For all five of these genes, we identified transcripts with a 3′ CCA tail, which is a typical feature of mature tRNAs. This tail is not genomically encoded in any of the genes and is presumably added post-transcriptionally by a tRNA nucleotidyltransferase (von Braun et al. 2007). Mature tRNAs also undergo extensive post-transcriptional base modifications (Suzuki 2021), some of which cause nucleotide misincorporations and deletions during reverse transcription (RT) (Clark et al. 2016; Behrens et al. 2021; Ceriotti et al. 2024). In the two fungal-derived tRNAs with suitable read depth for analysis, we found an increase in sequence variants around position 37 (Figure 1c), which is immediately 3′ of the anticodon and a known site for base modifications in many tRNAs (Schweizer et al. 2017). However, the overall misincorporation rate for both of these tRNAs was low, including at sites such as 26G that show signatures in many (but not all) plant mt-tRNAs (Ceriotti et al. 2024). The sequence variants at the 5′ and 3′ ends of tRNA-Met (Figure 1c) are likely mapping artefacts resulting from variation in transcript length. We found that many tRNA-Met reads are tailed with the post-transcriptional addition of CA instead of CCA. Because the preceding base (i.e., the discriminator base) is a C, both modifications result in a CCA tail but with different endpoints relative to the reference gene model. RT-mediated terminal transferase activity at the 5′ end of these tRNA-Met transcripts may also lead to artefactual mismatches relative to the reference sequence (Figure 1c).

The fungal-derived tRNA-Gln and tRNA-Met transcripts retained intact CCA tails after periodate treatment at levels similar to other mitochondrial tRNAs (Figure 1d), indicating that their 3′ ends are protected by an amino acid. For tRNA-Met, the retention of the 3′ nucleotide was much less common (2.6-fold lower on average) for transcripts that carried a post-transcriptional CA addition instead of CCA (see above), suggesting that the full-length tRNA-Met sequence is important for aminoacylation (Table S2). Inclusion of a synthetic spike-in control tRNA confirmed that the periodate treatment was effective in removing the 3′ nucleotide of uncharged tRNAs (Figure 1d). Although precise estimates of aminoacylation levels for the other fungal-derived tRNAs are not feasible due to their extremely low read abundance, the small number of detected transcripts exhibited little or no retention of the 3′ nucleotide after periodate treatment, suggesting that they are generally not aminoacylated.

The fact that tRNA-Gln is expressed and aminoacylated is notable because this gene is right in the middle of the initial fungal HGT that preceded the divergence of the Orchidaceae (Sinn and Barrett 2020), possibly explaining why this insertion has been subsequently retained. In contrast, the fungal-derived tRNA-Met gene is part of the larger secondary transfer, which also contains six protein-coding genes. Because none of these six genes retain an intact reading frame or exhibit mRNA-seq coverage above background levels for intergenic regions in the *C. maculata* mitogenome (Figures 2 and S2), the expressed tRNAs may be the sole functional component of the fungal-derived HGTs.

**Figure 2.**
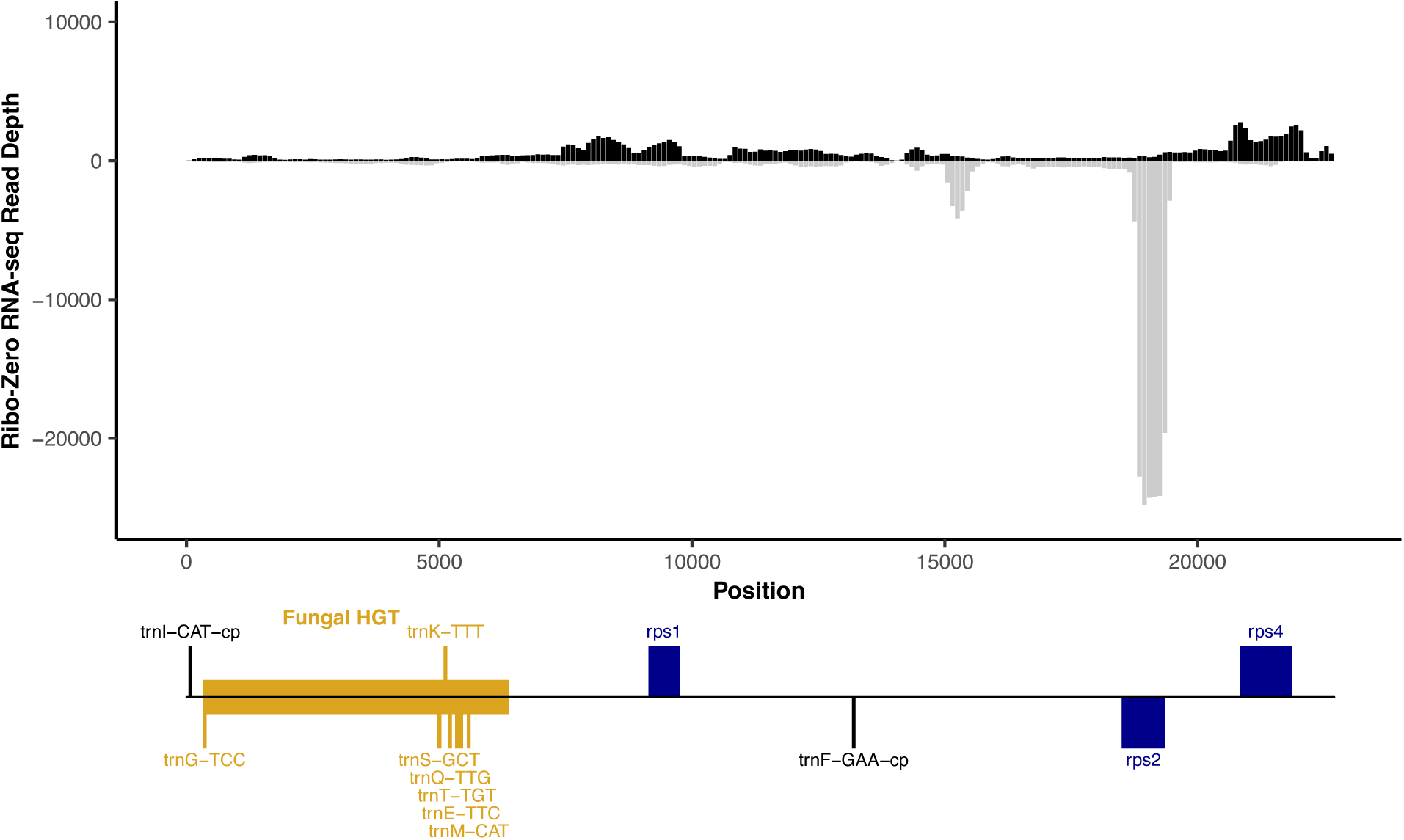
No evidence for functional expression of fungal-derived protein-coding genes in the *Corallorhiza maculata* mitochondrial genome. Transcript abundance was measured from rRNA-depleted (Ribo-Zero) RNA-seq mapped to the mitochondrial scaffold containing the fungal HGT region (Scaffold 8) from an assembly of total cellular DNA (see Figure S2 for mapping to other mitochondrial scaffolds). RNA and DNA were collected from the same individual (Plant 1). Depth is measured as average read sequence coverage (read count per position) for 100-bp windows across the length of the scaffold. Positive (black) and negative (gray) values indicate transcripts expressed from the forward and reverse orientations of the reference sequence, respectively. Mapping of sequencing reads from a second plant showed highly correlated read abundance across windows (*r* = 0.97 for log-transformed read depths). Annotated genes are indicated in the diagram below the x-axis. Genes above and below the line correspond to being encoded on the forward and reverse strand of the reference sequence, respectively. The region that was horizontally acquired from fungi and associated tRNA genes are indicated in gold. In addition to the annotated tRNA genes in this region, it contains apparent pseudogenes derived from the fungal protein-coding genes *atp6*, *atp8*, *cox1*, *cox2*, *nad1*, and *nad4*. The tRNA genes outside this region on this scaffold are both plastid-derived. Because of column-based RNA purification, library size selection, and the challenges inherent in sequencing tRNAs, these RNA-seq libraries are not expected to capture mature tRNA expression. Note that transcript abundance across the fungal-derived region does not exceed background levels in intergenic region on this or other scaffolds (Figure S2).

Interestingly, neither of the two fungal-derived tRNAs that appear to be functionally expressed and aminoacylated expand the decoding capacity of the *C. maculata* mitogenome because native mitochondrial and/or plastid-derived tRNAs with the same anticodons are also present. Instead, the *C. maculata* mitogenome offers a striking example of tRNAs from diverse evolutionary origins being redundantly expressed in the same organelle (Figure 1b). This redundancy may represent an intermediate state in a more general evolutionary process responsible for the exceptional propensity of plant mitogenomes for tRNA HGT and eventual functional replacement (Small et al. 1999; Warren and Sloan 2020; Warren, Salinas-Giegé, Triant, et al. 2021).

### Corallorhiza maculata maintains typical plant enzymatic machinery for aminoacylation of organellar tRNA-Gln and tRNA-Met despite tRNA HGT

The expression and apparent aminoacylation of the fungal-derived tRNA-Gln and tRNA-Met raise questions about the identity of the aaRSs charging these HGT tRNAs. Plant nuclear genomes typically encode two separate sets of aaRSs for aminoacylating cytosolic vs. organellar tRNAs (Duchêne et al. 2005). In *C. maculata*, we identified transcripts coding for typical cytosolic and organellar MetRS enzymes, and only the organellar MetRS was predicted to be targeted to the mitochondria (Table S3). In contrast, we did not find any MetRS sequences of fungal origin in the *C. maculata* transcriptome. Both findings suggest that a typical plant organellar MetRS is the sole enzyme for charging tRNA-Met in *C. maculata* mitochondria.

Aminoacylation of tRNA-Gln in plant and fungal mitochondria occurs via an indirect pathway in which the tRNA is first charged with Glu by a non-discriminating GluRS, and then Glu is converted to Gln by a tRNA-dependent amidotransferase (GatCAB in plant mitochondria and GatFAB in fungal mitochondria) (Pujol et al. 2008; Frechin et al. 2009; Araiso et al. 2014). All three subunits of the plant GatCAB complex were detected in the *C. maculata* transcriptome with predicted targeting to the organelles (Table S3), whereas no fungal-type GatFAB subunits were found. Additionally, we did not detect any fungal GlnRS/GluRS HGT or retargeting of the *C. maculata* cytosolic GlnRS enzyme to mitochondria (Table S3). These results suggest that the typical plant transamidation pathway is the only mechanism for charging tRNA-Gln in *C. maculata* mitochondria. However, we cannot confidently conclude that the HGT tRNAs are “correctly” loaded with the cognate amino acid, because the tRNA-seq method used in this study can only differentiate between charged and uncharged tRNAs and does not identify which amino acids are loaded onto the tRNAs. Understanding how these foreign tRNAs interact with plant enzymatic machinery despite more than a billion years of sequence divergence from their native tRNA counterparts is a fascinating area for future investigation.

## Methods

### Tissue collection and RNA/DNA extraction

Shoot tissue was collected from two *C. maculata* individuals growing <10 m apart near Fort Collins, CO, USA (40° 33′ 49″ N, 105° 11′ 0″ W). Samples were collected at 11:00am on June 4, 2024 and stored on ice until 1:30pm when RNA extraction was performed using an acid-phenol method as previously described (Ceriotti et al. 2024). Two extractions were performed per plant, each using a whole flower sampled from near the shoot apex. An additional flower from each plant was used for DNA extraction (Qiagen DNeasy kit).

### Organelle genome sequencing and analysis

Library construction and sequencing of *C. maculata* genomic DNA samples were performed by Novogene, using an NEBNext Ultra II DNA Library Prep Kit and a 2×150 bp run on an Illumina NovaSeq X platform. A subset of 15M read pairs from the “Plant 1” sample was trimmed with Cutadapt v4.0 (Martin 2011) and assembled with SPAdes v4.0.0, using the meta option (Nurk et al. 2017). Putative mitogenome-derived contigs were extracted based on coverage depth (40-150×) and length (>1000 bp) and then further screened for the presence of known plant mitochondrial gene sequences, resulting in a set of 23 contigs with a total length of 495,364 bp. To assemble a reference plastome, NOVOPlasty v4.3.1 (Dierckxsens et al. 2017) was seeded with the *accD* gene sequence from GenBank accession KM390016.1 (Barrett et al. 2014) and used to analyze the full set of trimmed reads from the Plant 1 sample. Organelle tRNA genes were identified with tRNAscan-SE 2.0 (Chan and Lowe 2019) followed by manual curation.

### Charged tRNA-seq

To measure tRNA expression and infer aminoacylation states based on retention of CCA tails, an MSR-seq protocol (Watkins et al. 2022) was performed on two biological replicates from each of two *C. maculata* plants both with and without periodate treatment. Library construction, sequencing, and data analysis (including quantification of read abundance, CCA-tailing, and base misincorporations) were performed as described previously (Ceriotti et al. 2024). The reference tRNA set for mapping included annotated tRNAs from *C. maculata* organellar genomes (see above), *Arabidopsis thaliana* nuclear tRNAs from the PlantRNA 2.0 database (Cognat et al. 2022), and a *Bacillus subtilis* tRNA-Ile that was synthesized and used as an internal “spike-in” control as described previously (Ceriotti et al. 2024).

### rRNA-depleted RNA-seq

RNA samples (see above) from a single biological replicate for each *C. maculata* plant (samples 1A and 2A) were processed with a Qiagen RNeasy MinElute kit to remove contaminating genomic DNA and then used to generate strand-specific rRNA-depleted RNA-seq libraries (Illumina TruSeq Stranded Total RNA with Ribo-Zero Plant kit) and sequenced on a 2×150 bp run of an Illumina NovaSeq X platform. Library construction and sequencing was performed by Novogene. The resulting reads were trimmed with Cutadapt and mapped to *C. maculata* mitogenome scaffolds with Bowtie2 v2.2.5 using default parameters (Langmead and Salzberg 2012), and resulting alignments were analyzed with Samtools v1.17 (Li et al. 2009) and custom scripts to quantify mRNA and long non-coding RNA transcript abundance. Trimmed reads were also assembled *de novo* with Trinity v2.15.2 (Grabherr et al. 2011) and searched for homologs of aaRSs and GatCAB/GatFAB with NCBI TBLASTN v2.14.1+ (Camacho et al. 2009). Subcellular localization was predicted with TargetP v2.0 (Emanuelsson et al. 2000).

## Data Availability

All raw sequencing reads are available via NCBI SRA (BioProject PRJNA1186809). Code and processed data files, including tRNA read counts, are available via GitHub (https://github.com/dbsloan/Corallorhiza_tRNAs). Transcriptome and genomic DNA assemblies are available via Zenodo (DOI: 10.5281/zenodo.14172377; https://zenodo.org/records/14172377).

## Acknowledgements

We thank Kate Wilsterman for assistance with collecting *C. maculata* tissue samples. This work was supported by funding from the National Science Foundation (MCB-2322154), an IUBMB Wood-Whelan Research Fellowship, and an HHMI Hanna H. Gray Fellowship.

## Supplementary Material

**Table S1.**
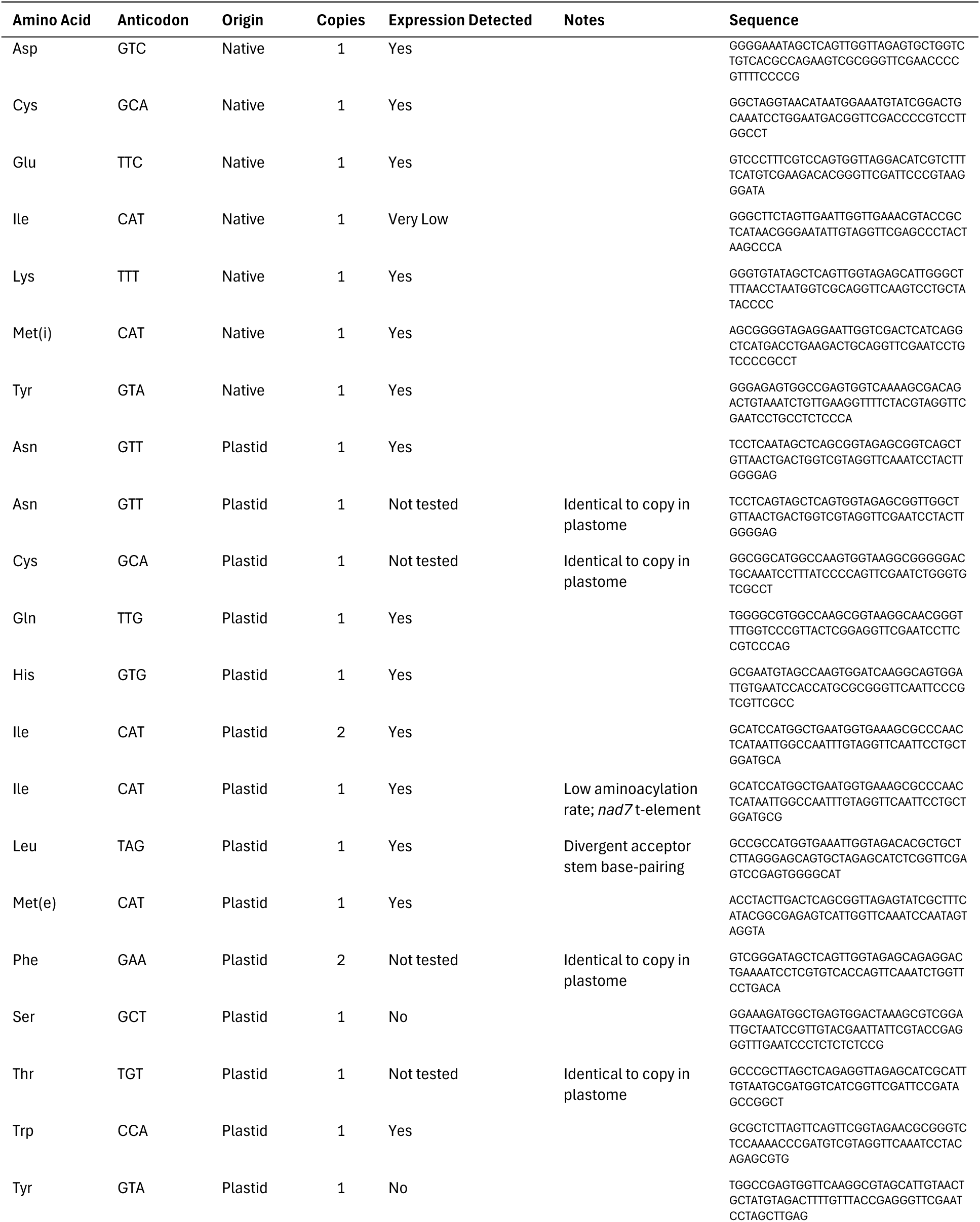
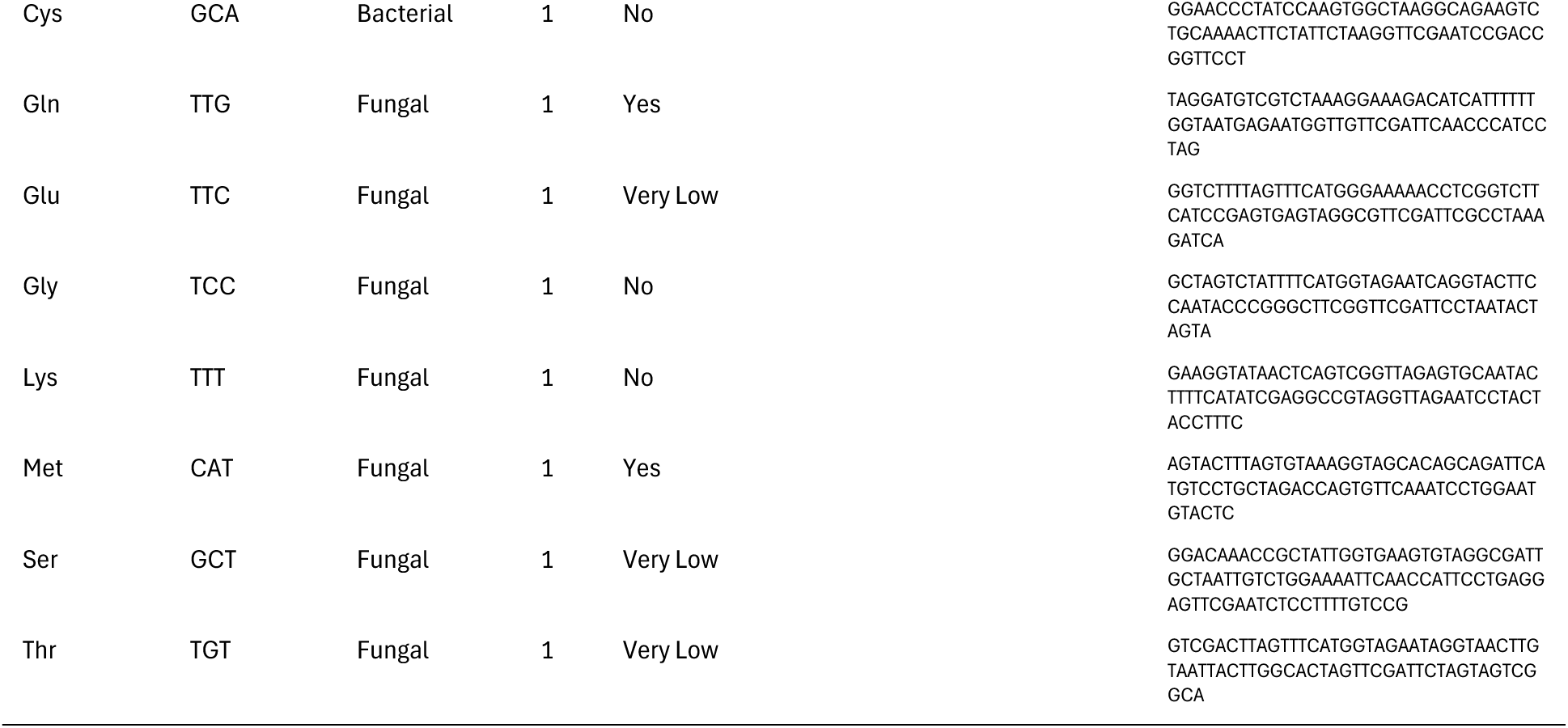
Summary of mitochondrial tRNA genes in *Corallorhiza maculata*

**Table S2.** Differential retention of 3′ terminal nucleotide following periodate treatment in fungal-derived tRNA-Met transcripts with CCA vs CA tails.

| Replicate | Proportion of Reads<br>(Control Libraries) |  | Proportion of Reads<br>(Periodate Libraries) |  | Periodate/Control<br>Ratio |  |
| --- | --- | --- | --- | --- | --- | --- |
|  | CCA | CA | CCA | CA | CCA | CA |
| 1A | 0.106 | 0.488 | 0.096 | 0.166 | 0.909 | 0.340 |
| 1B | 0.117 | 0.422 | 0.092 | 0.141 | 0.788 | 0.334 |
| 2A | 0.122 | 0.412 | 0.134 | 0.128 | 1.100 | 0.311 |
| 2B | 0.055 | 0.158 | 0.044 | 0.063 | 0.809 | 0.400 |
| **Mean** | **0.100** | **0.370** | **0.092** | **0.125** | **0.901** | **0.346** |

**Table S3.** Targeting predictions for *Corallorhiza maculata* enzymes involved in charging of tRNA-Gln and tRNA-Met. Transcript and protein sequences are available via GitHub (https://github.com/dbsloan/Corallorhiza_tRNAs).

| Gene | Arabidopsis Ref | Trinity Contig | TargetP Analysis |  |  |
| --- | --- | --- | --- | --- | --- |
|  |  |  | Prediction | Mito Prob | Plastid Prob |
| GatA | AT3G25660 | TRINITY_DN17257_c0_g1_i12 | Plastid | 0.009 | 0.666 |
| GatB | AT1G48520 | TRINITY_DN6546_c1_g1_i3 | Mito | 0.913 | 0.019 |
| GatC | AT4G32915 | TRINITY_DN31742_c0_g1_i1 | Mito | 0.771 | 0.080 |
| GlnRS (cytosolic) | AT1G25350 | TRINITY_DN21878_c0_g5_i5 | None | 0.000 | 0.000 |
| GluRS (organellar) | AT5G64050 | TRINITY_DN17518_c0_g1_i5 | Plastid | 0.236 | 0.643 |
| GluRS (cytosolic) | AT5G26710 | TRINITY_DN951_c0_g1_i5 | None | 0.006 | 0.091 |
| MetRS (cytosolic) | AT4G13780 | TRINITY_DN72148_c1_g1_i1 | None | 0.000 | 0.000 |
| MetRS (organellar) | AT3G55400 | TRINITY_DN1188_c0_g1_i5 | Plastid | 0.323 | 0.595 |

**Figure S1.**
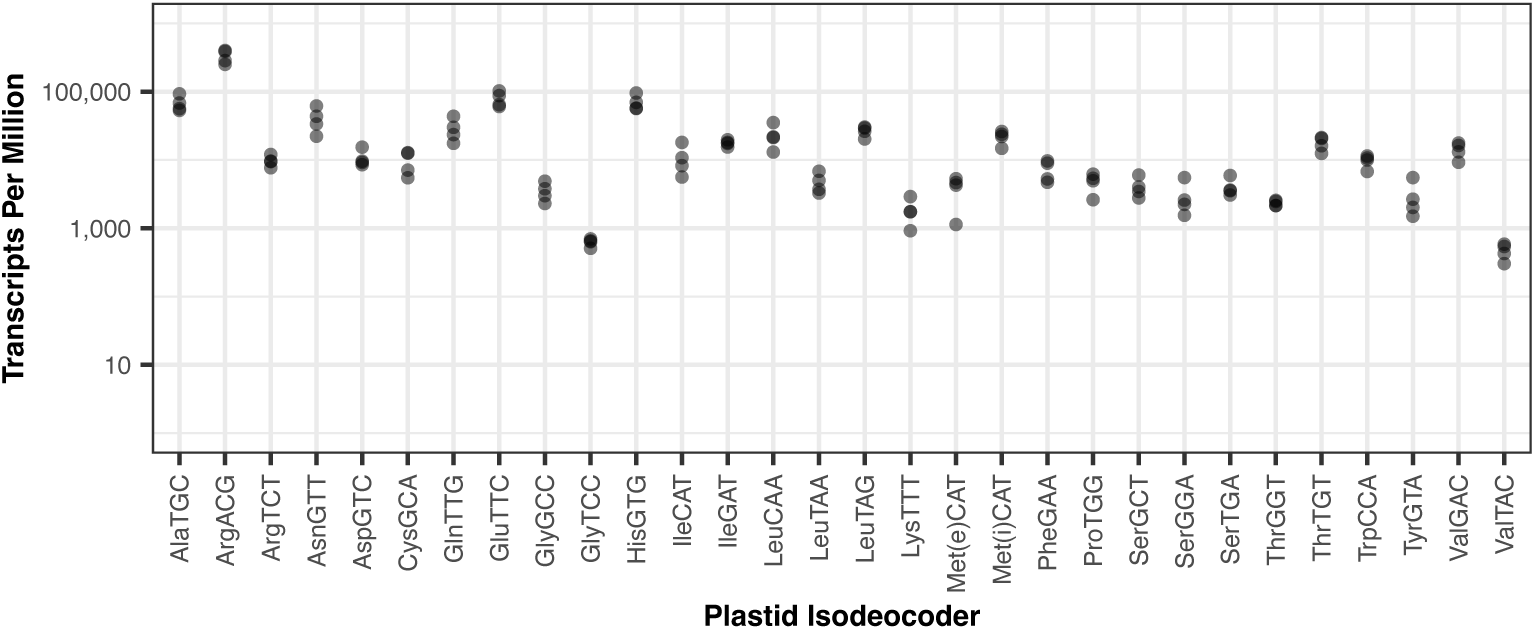
Expression of plastid-encoded tRNAs. Read abundances for each of four *Corallorhiza maculata* biological replicates (two flowers from each of two plants) are quantified as transcripts (reads) per million mapped to organellar reference sequences. Data are only plotted from control (no periodate) libraries and for reads that mapped uniquely to the reference. Note that read abundance for AsnGTT, CysGCA, PheGAA, and ThrTGT could also reflect expression of mitochondrial copies because there are identical copies of these plastid genes inserted in the mitogenome.

**Figure S2.**
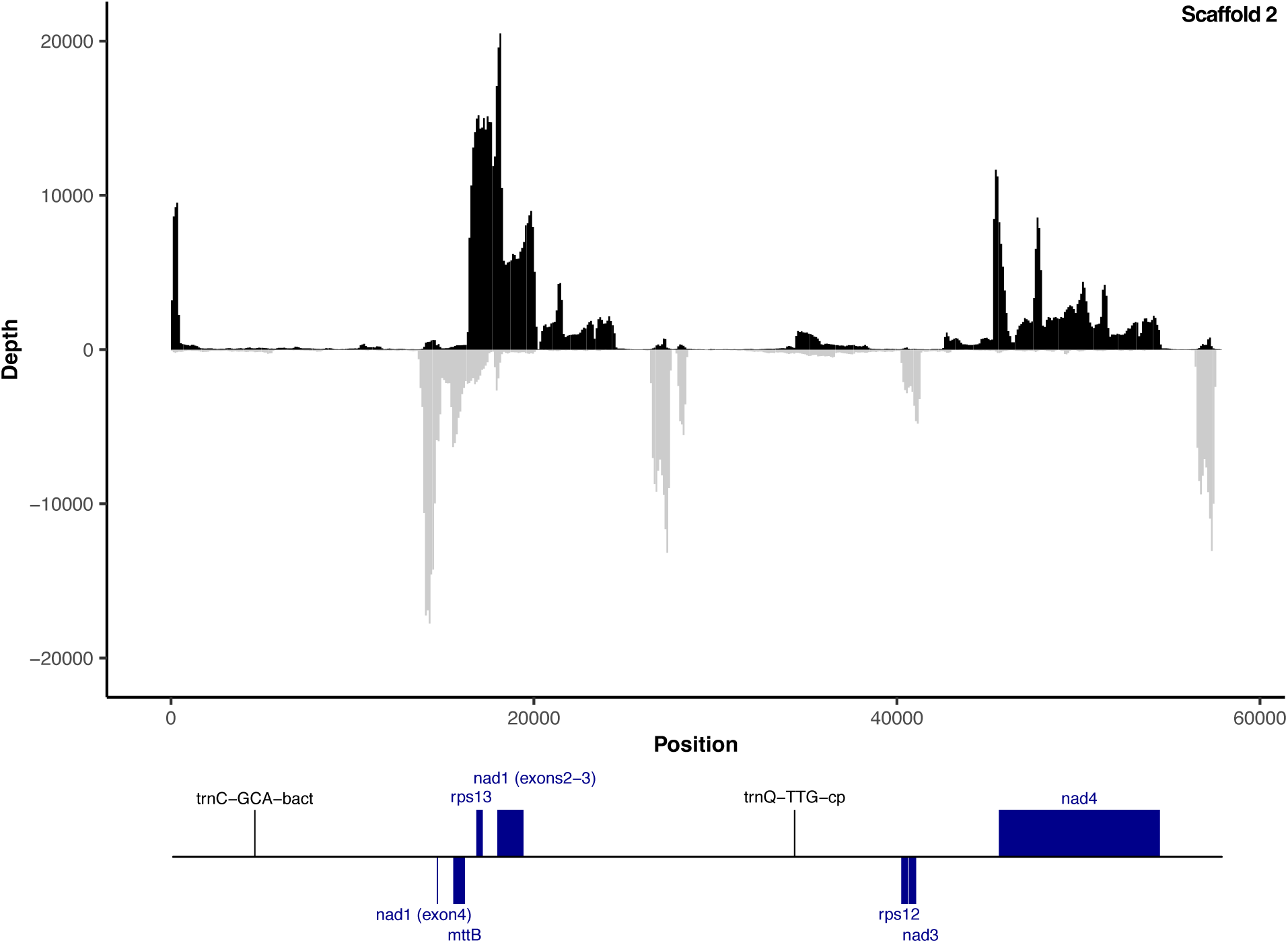
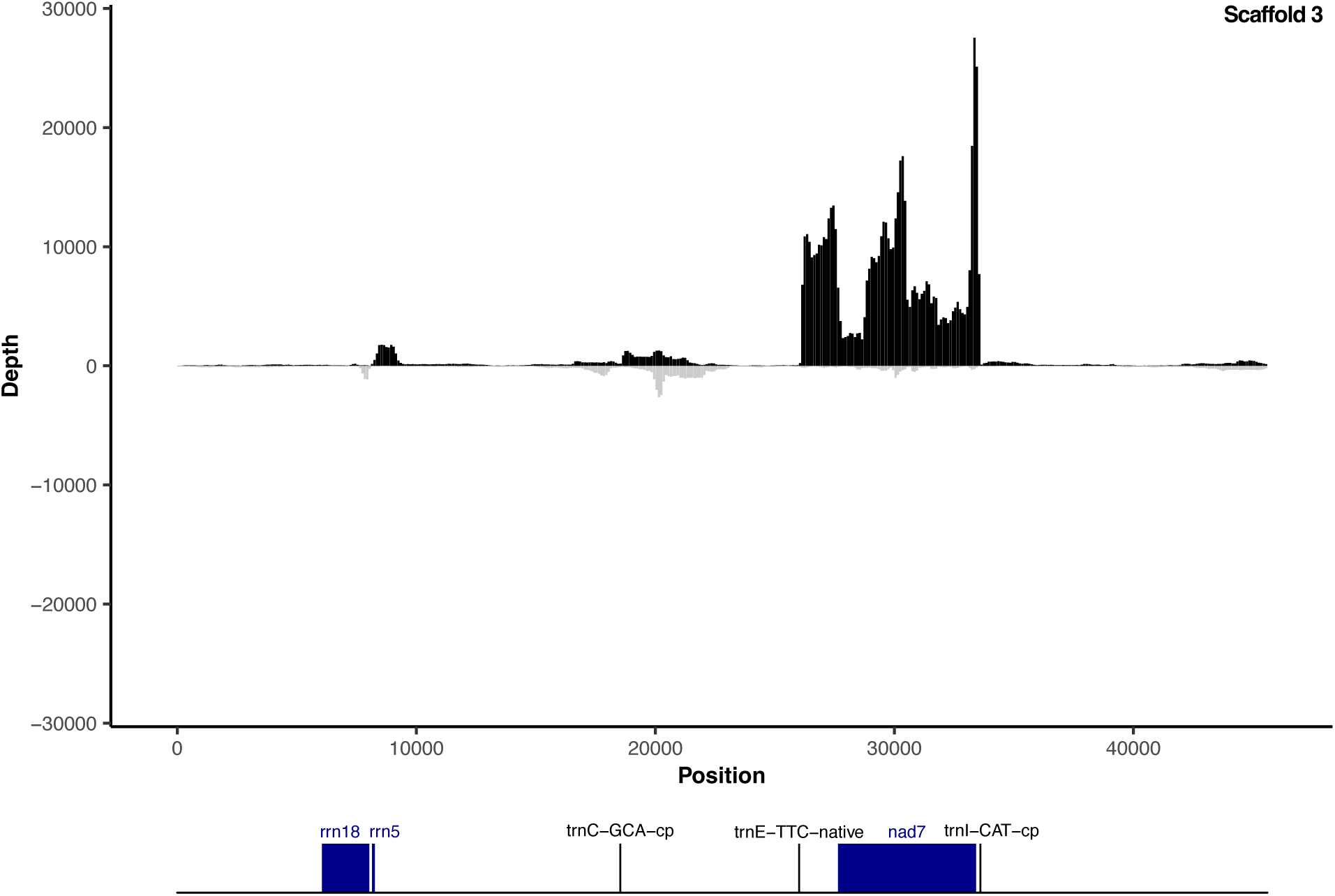
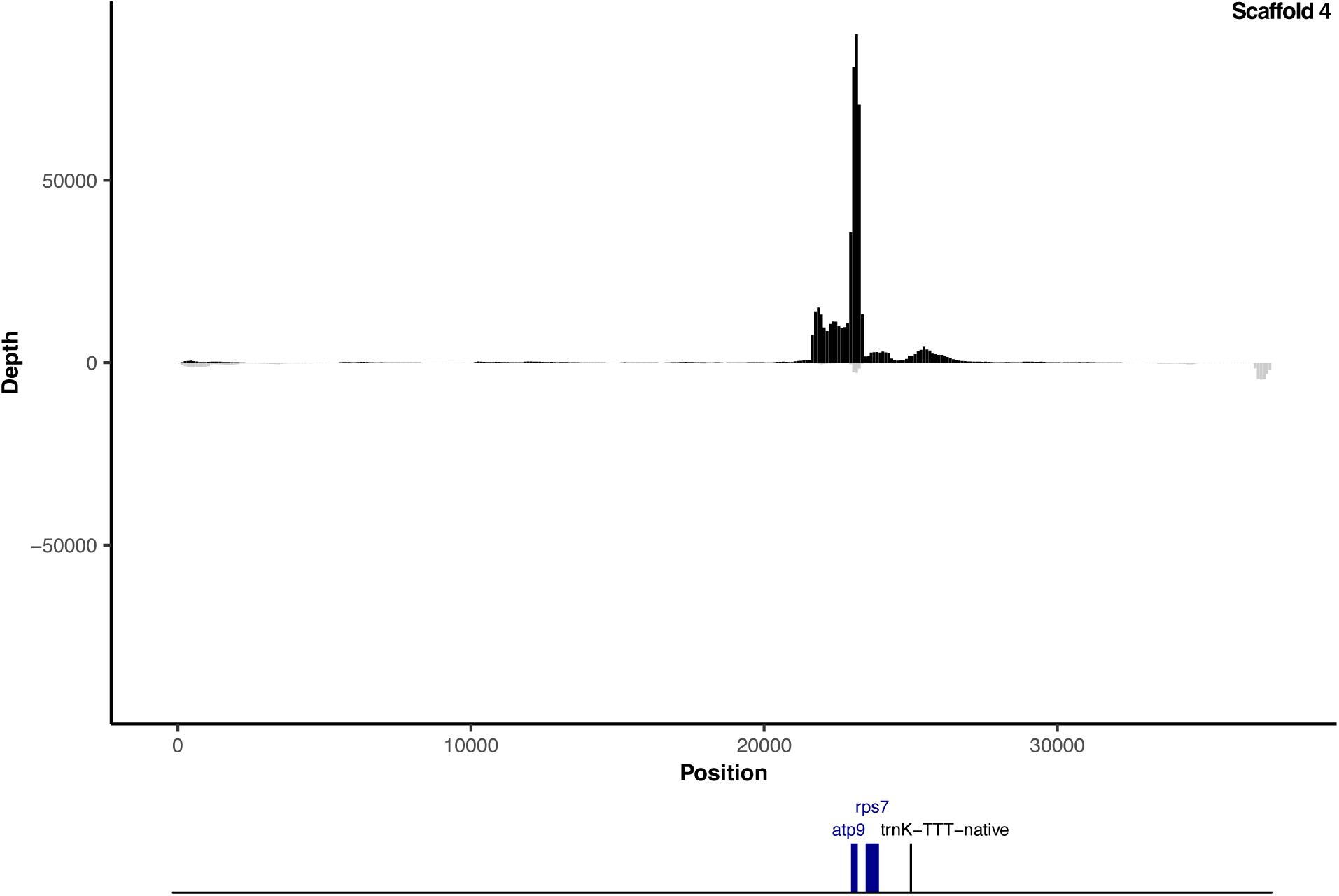
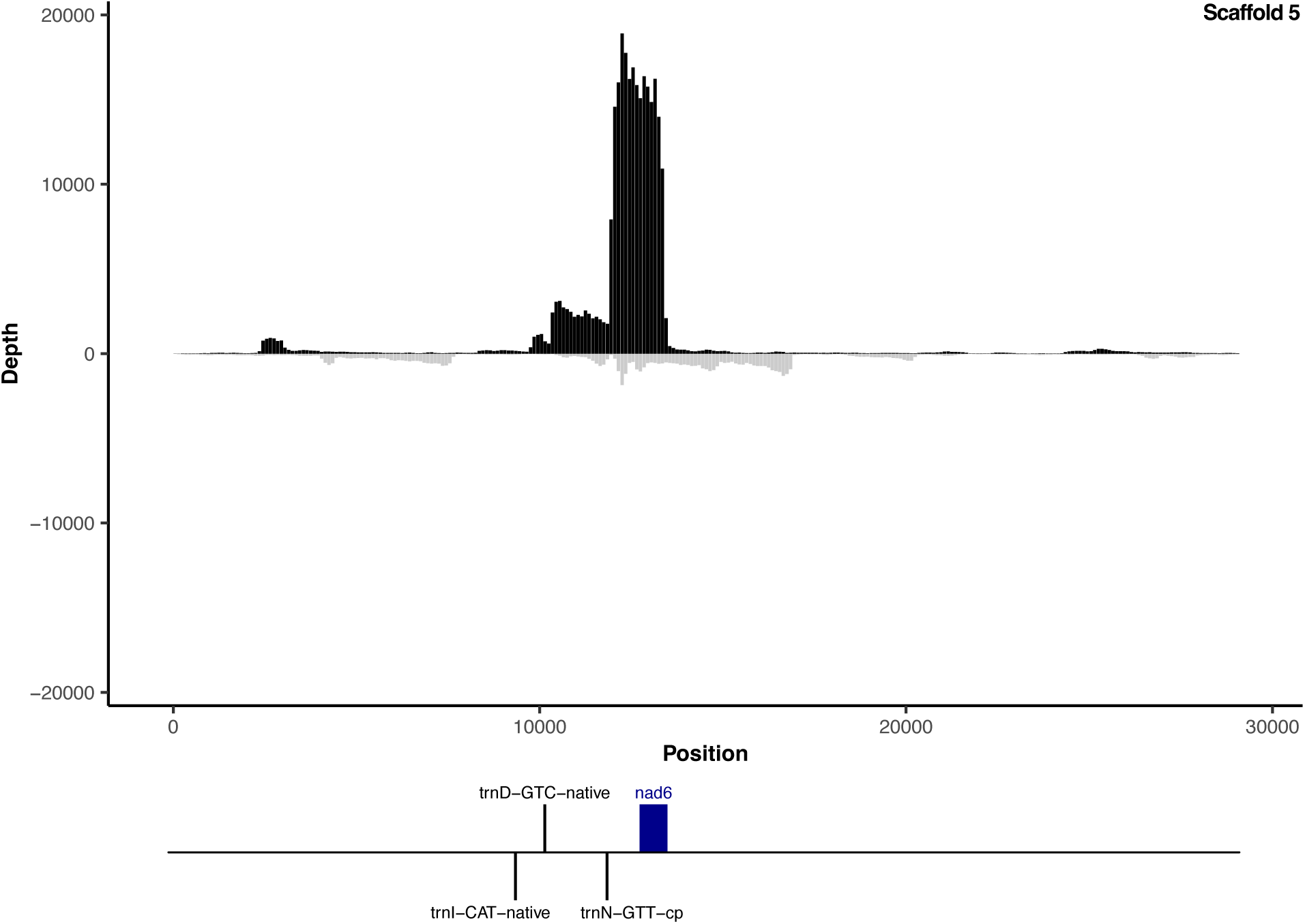
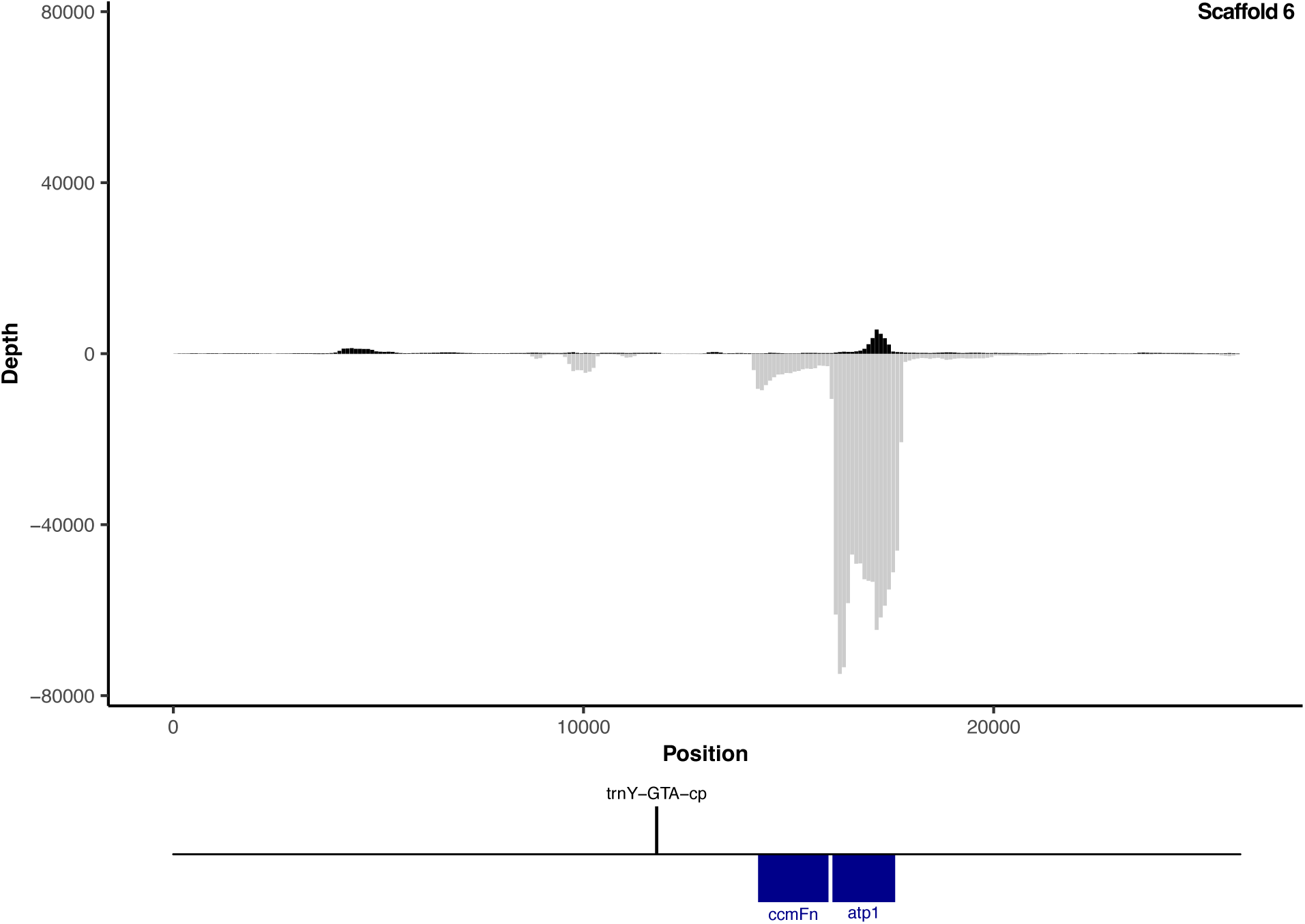
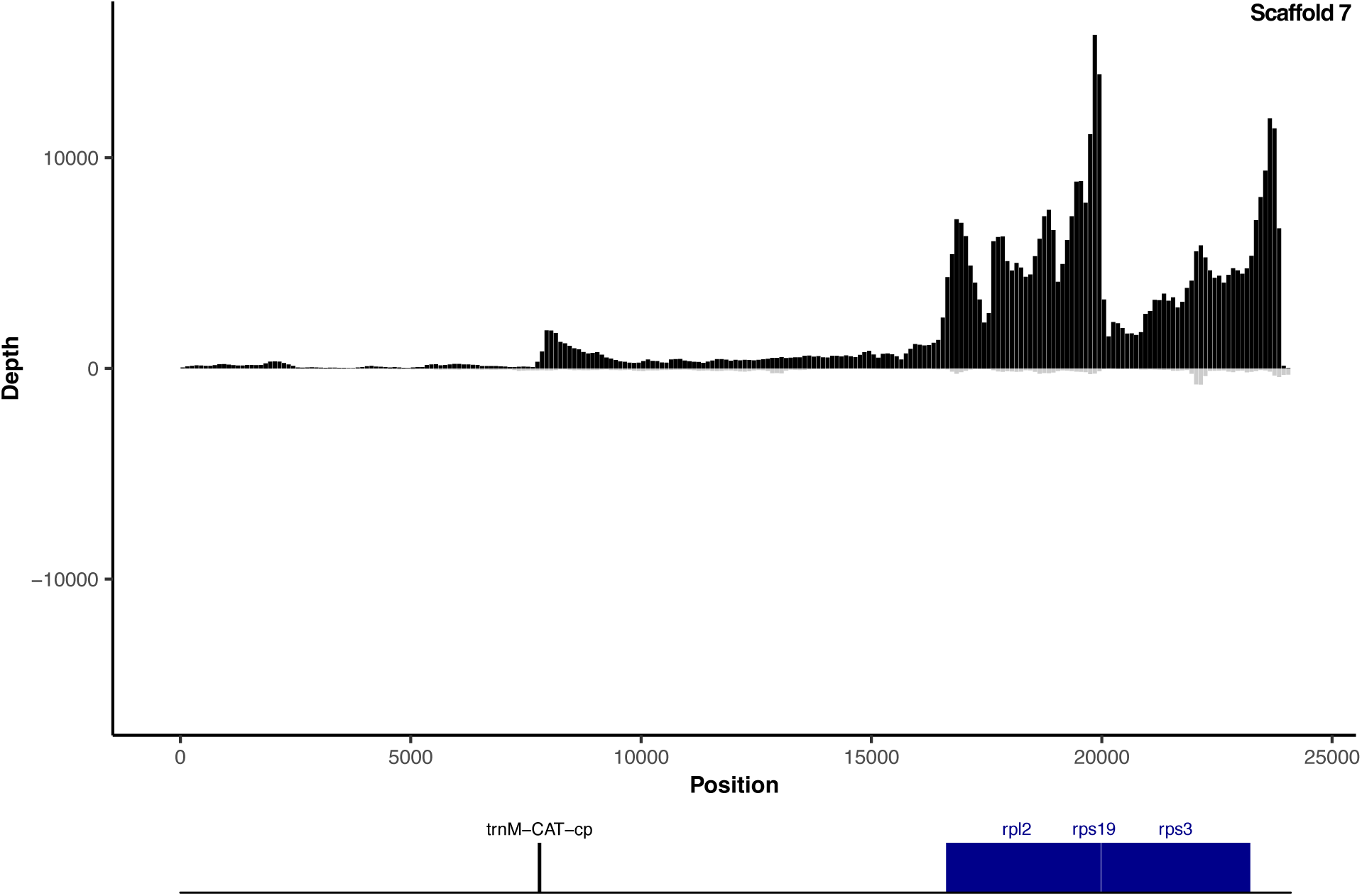
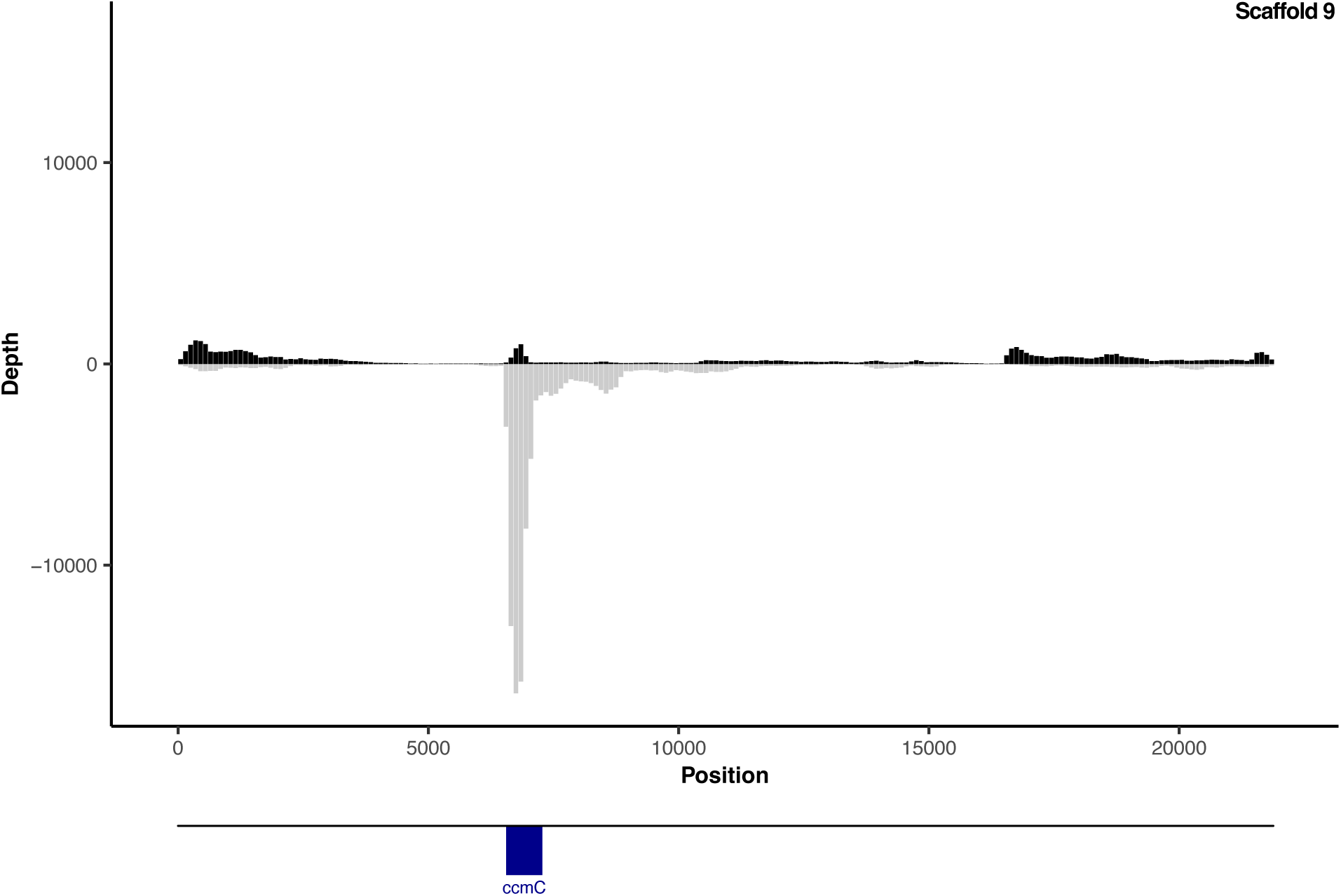
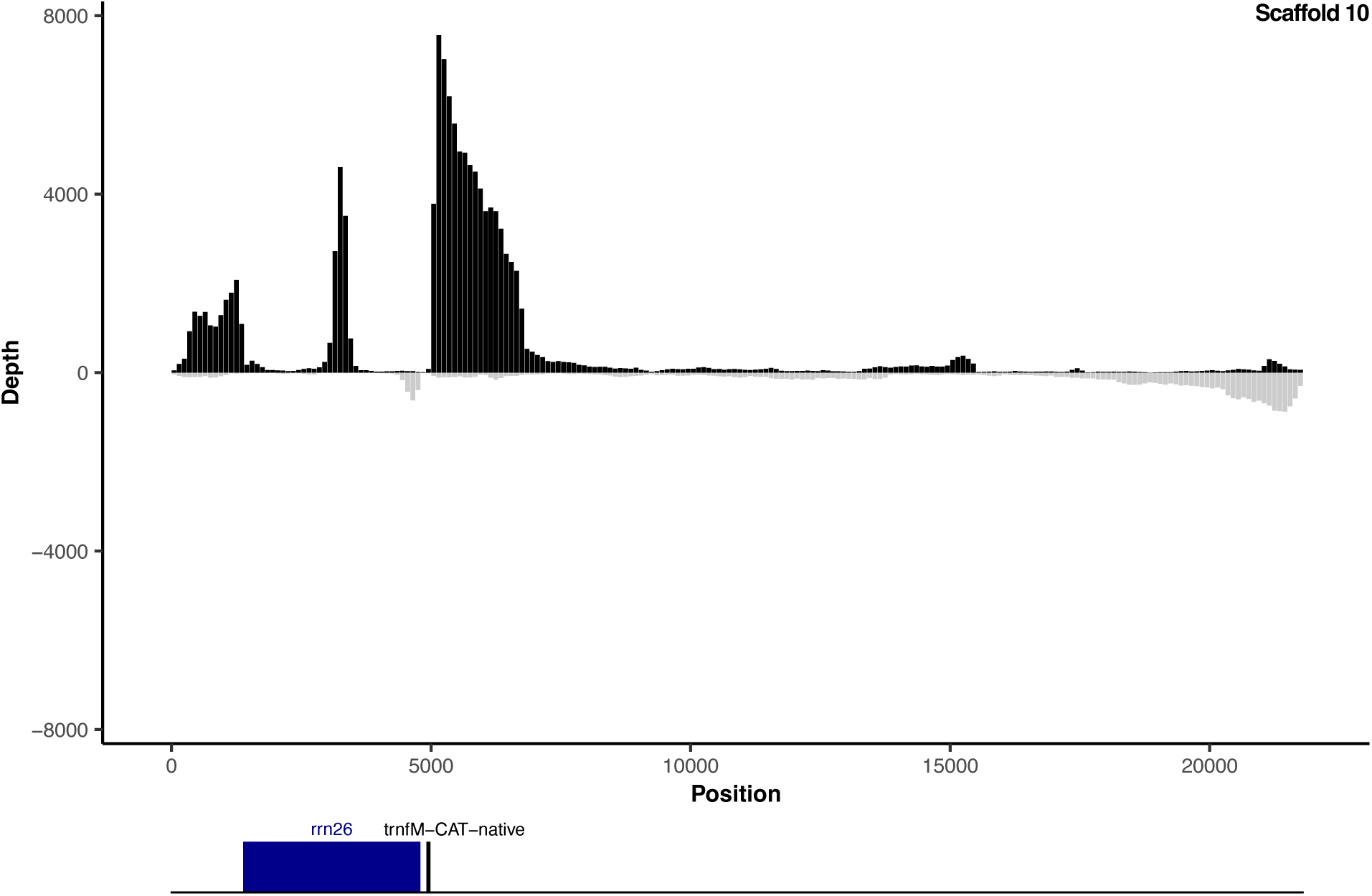
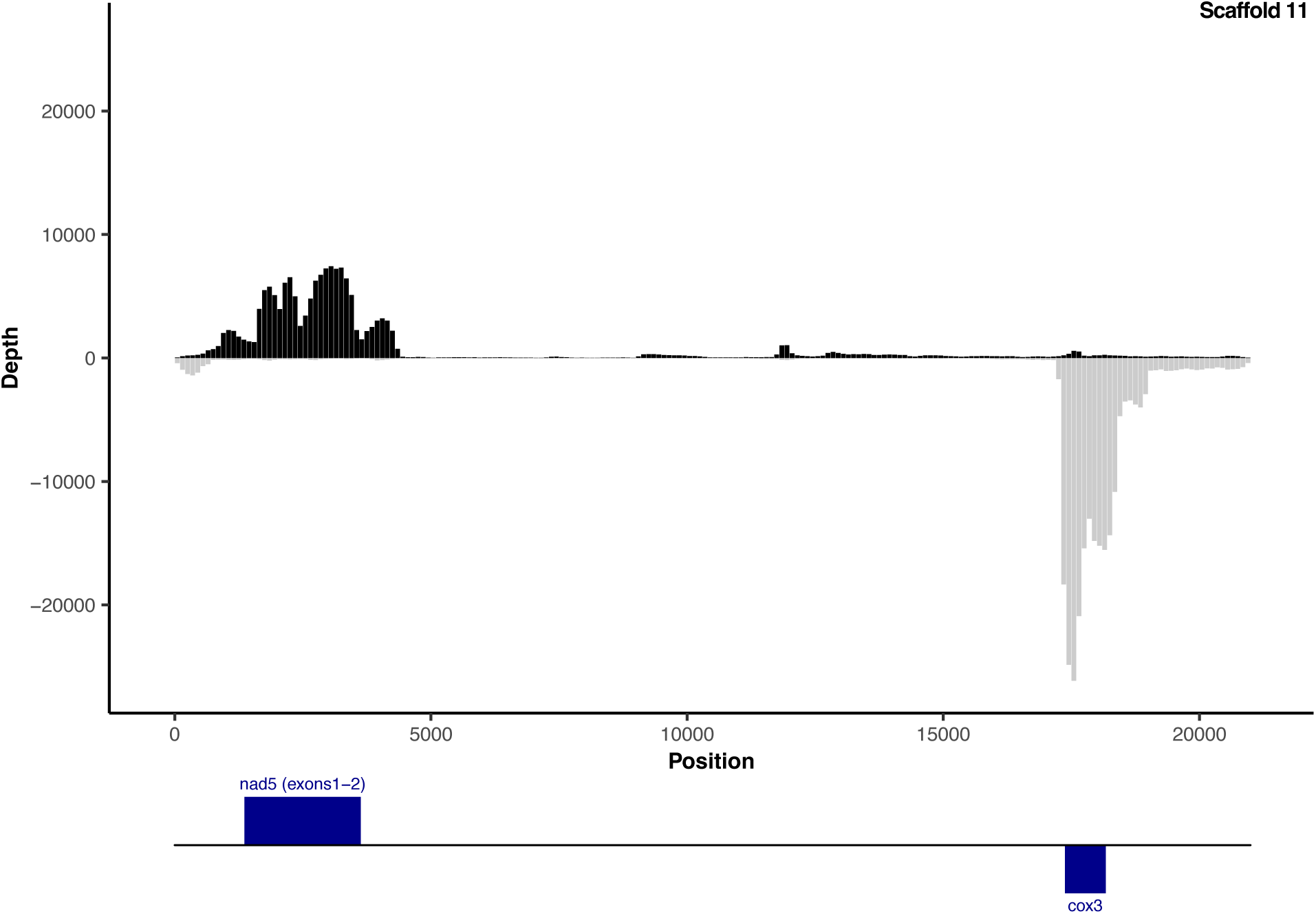
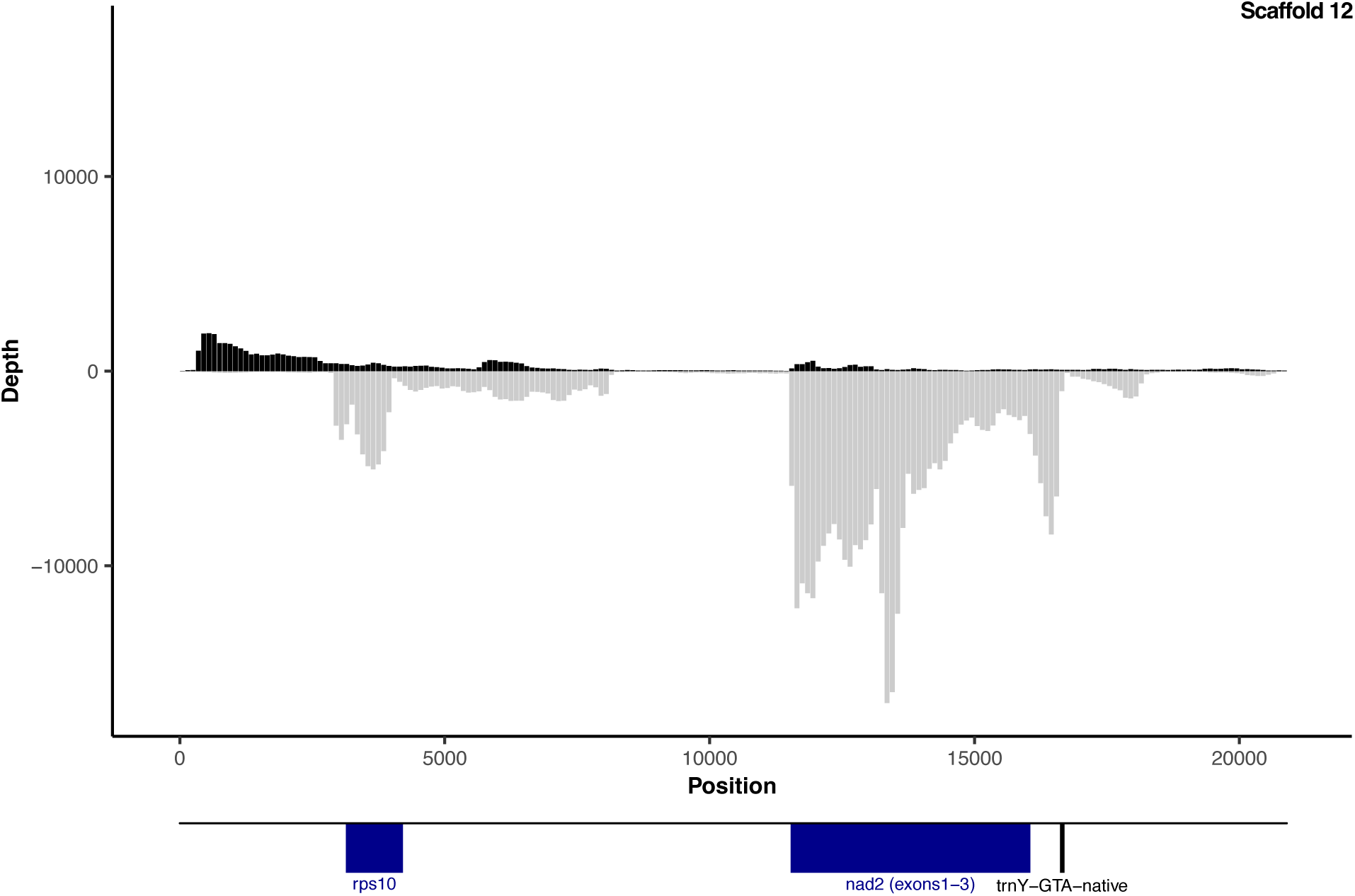
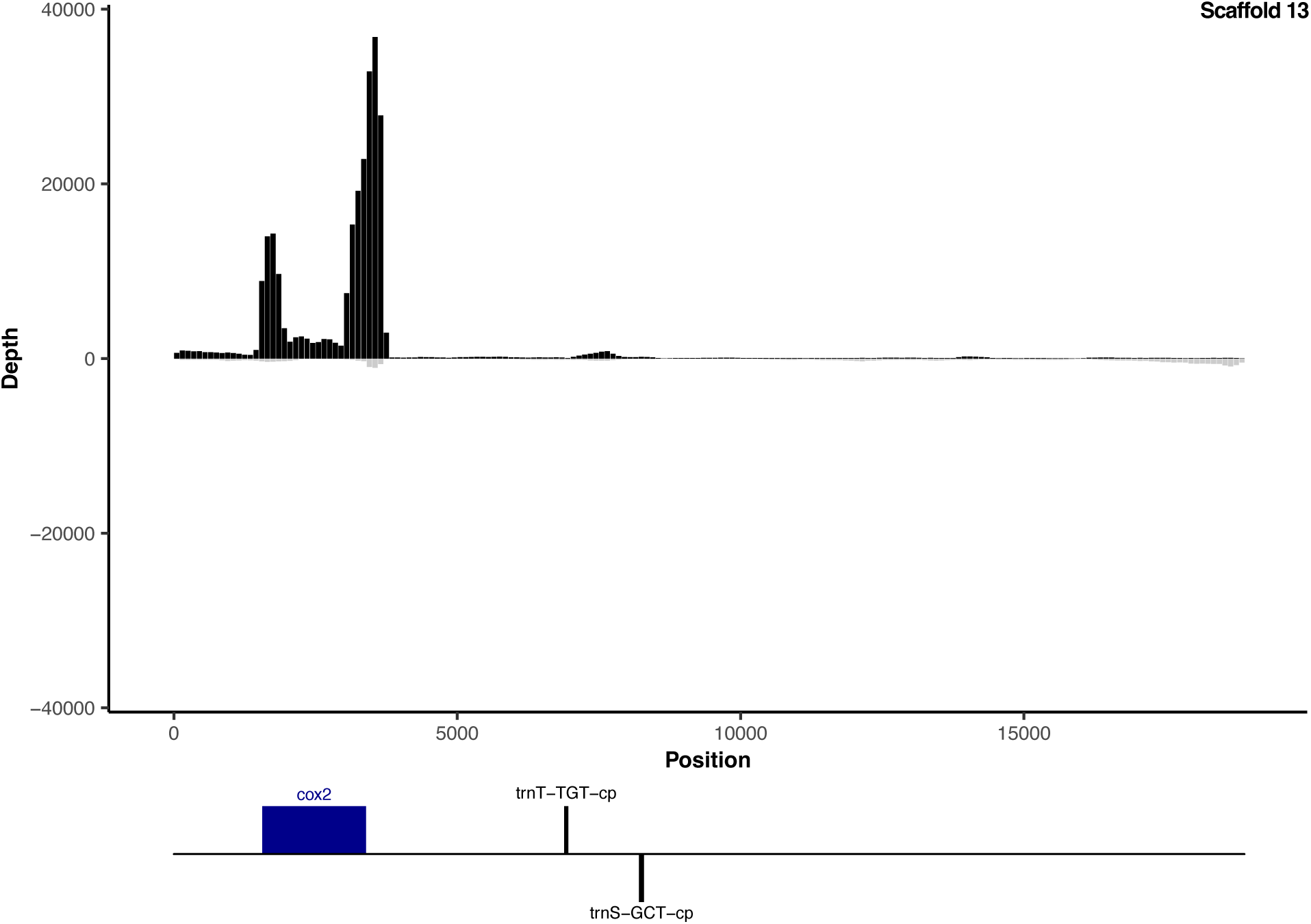
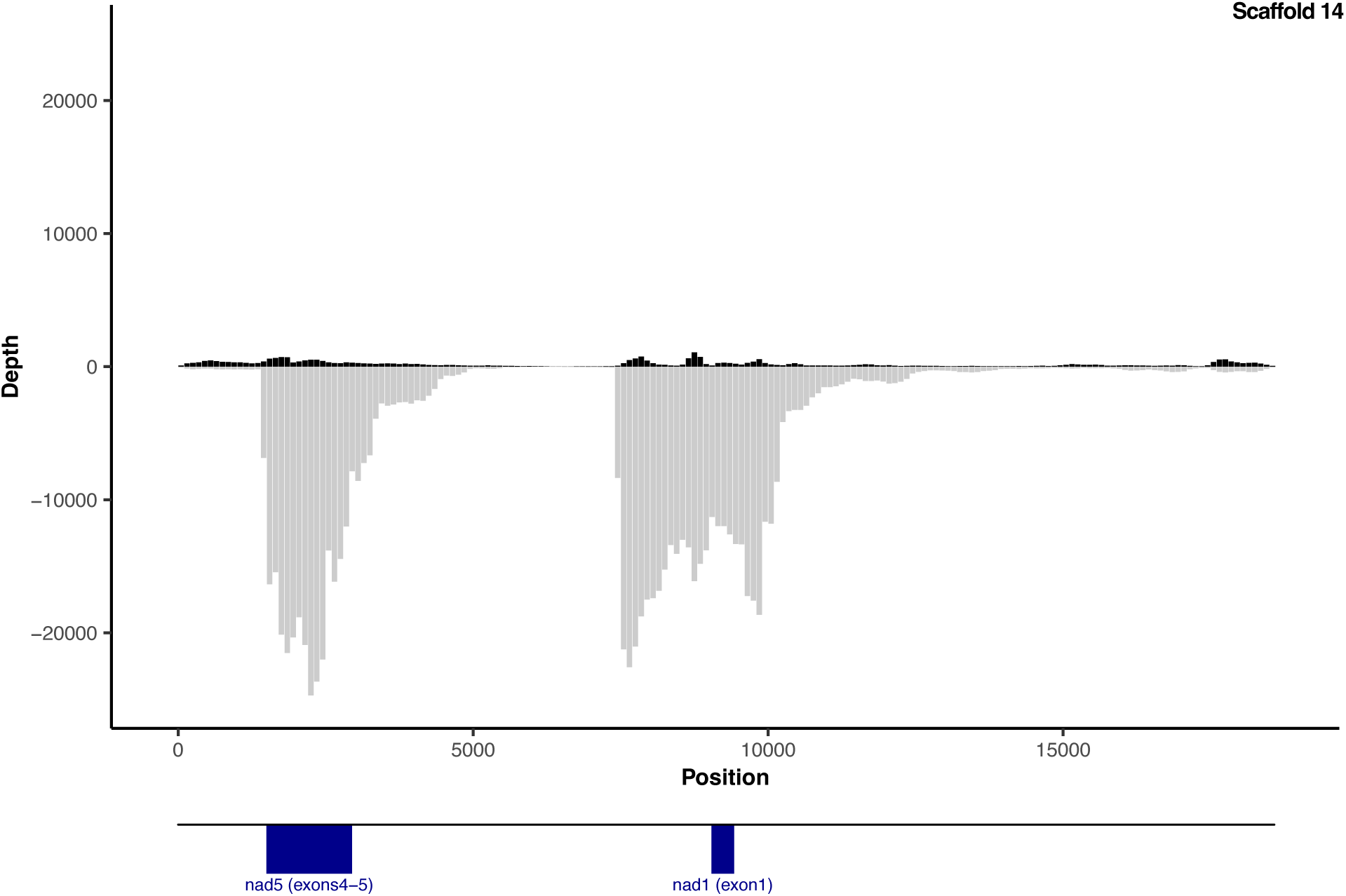
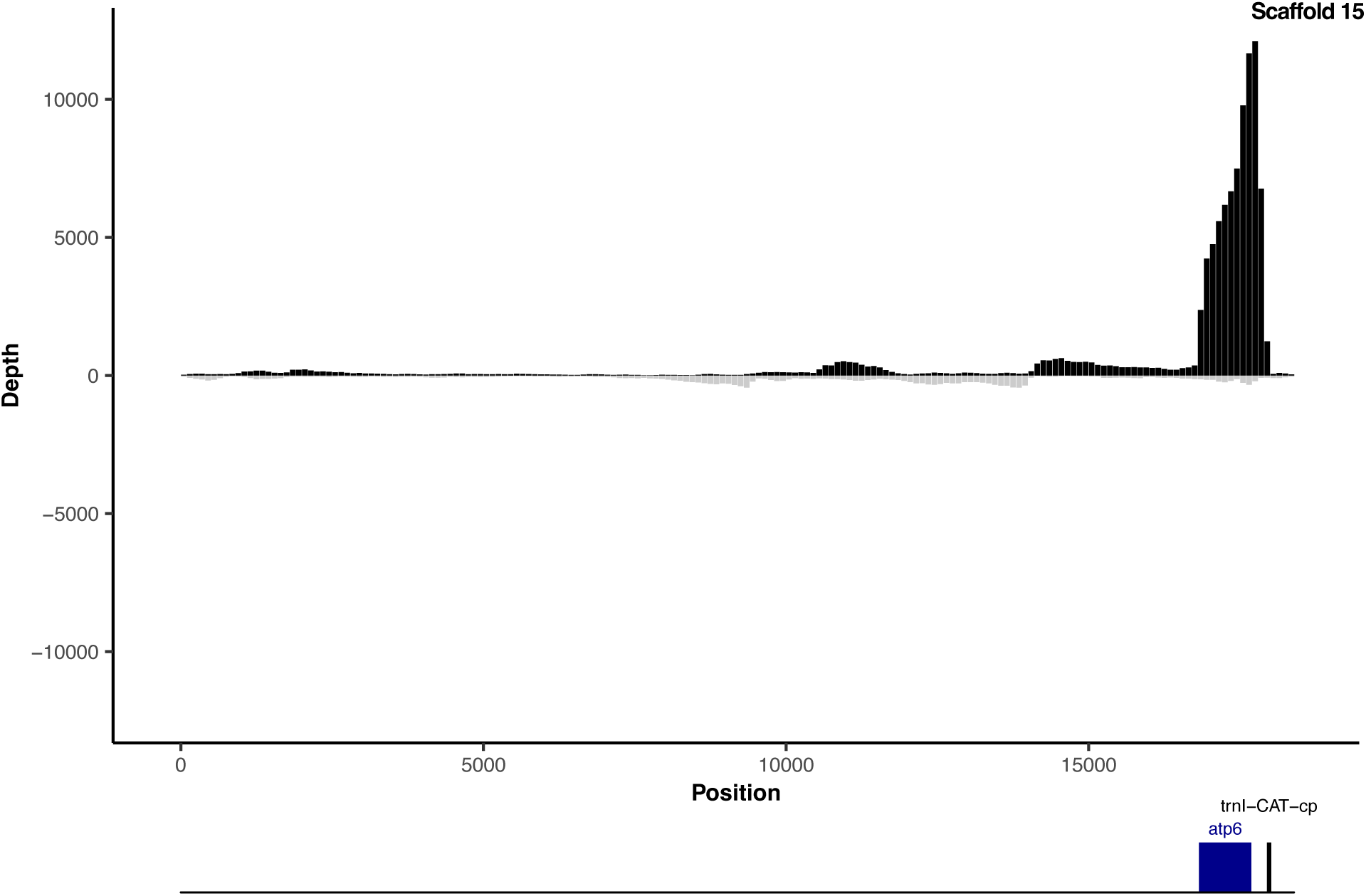
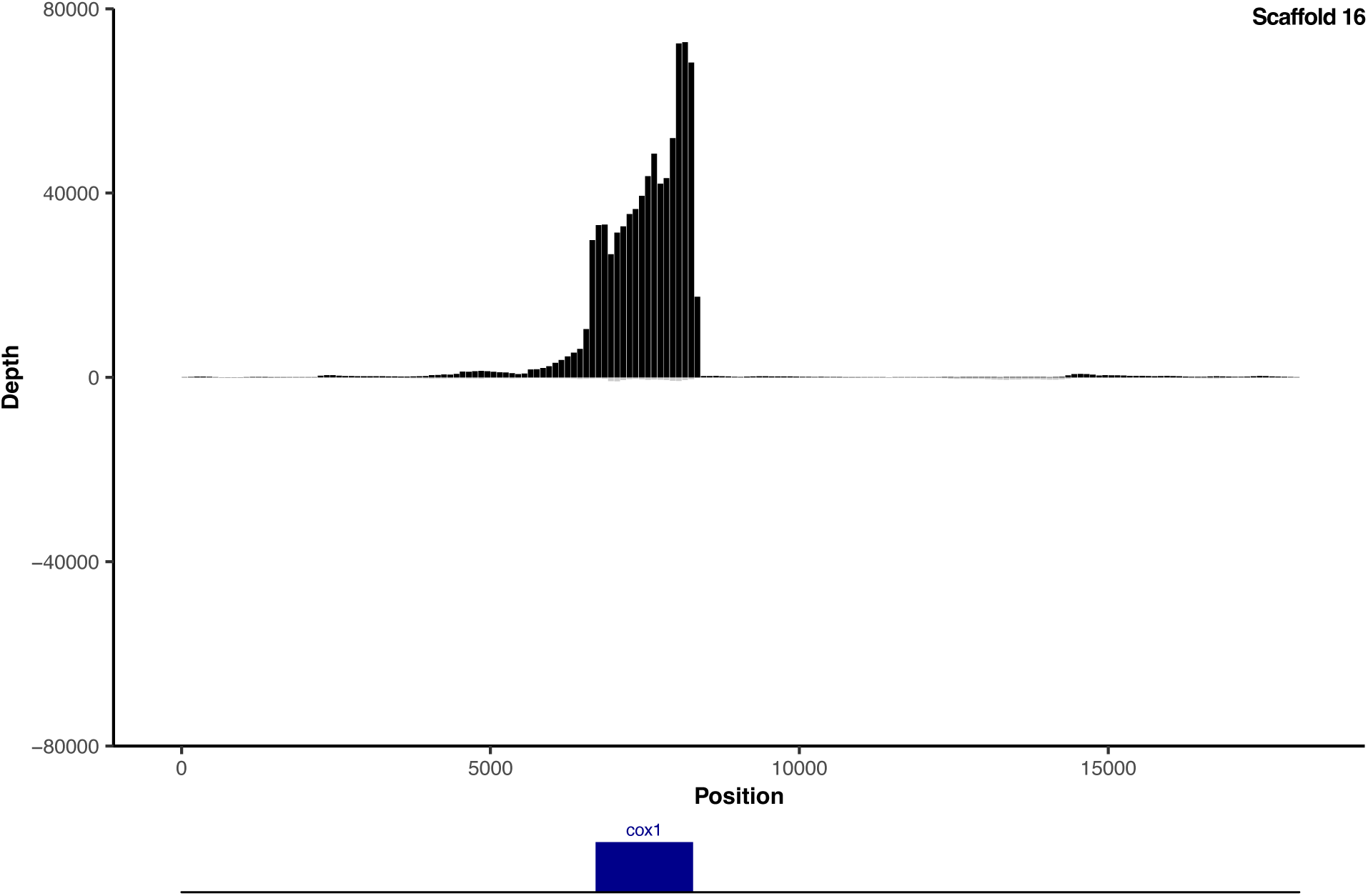
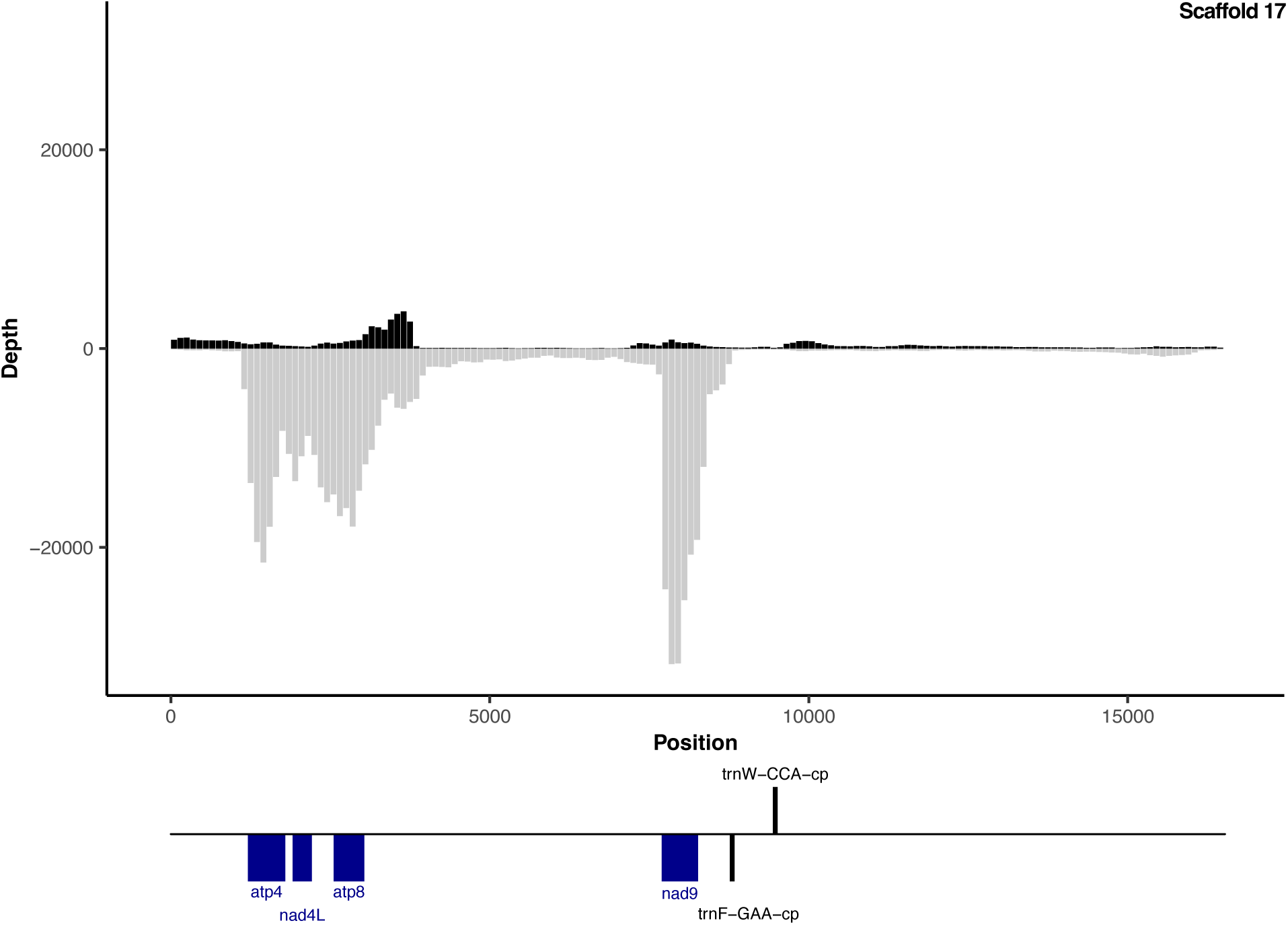
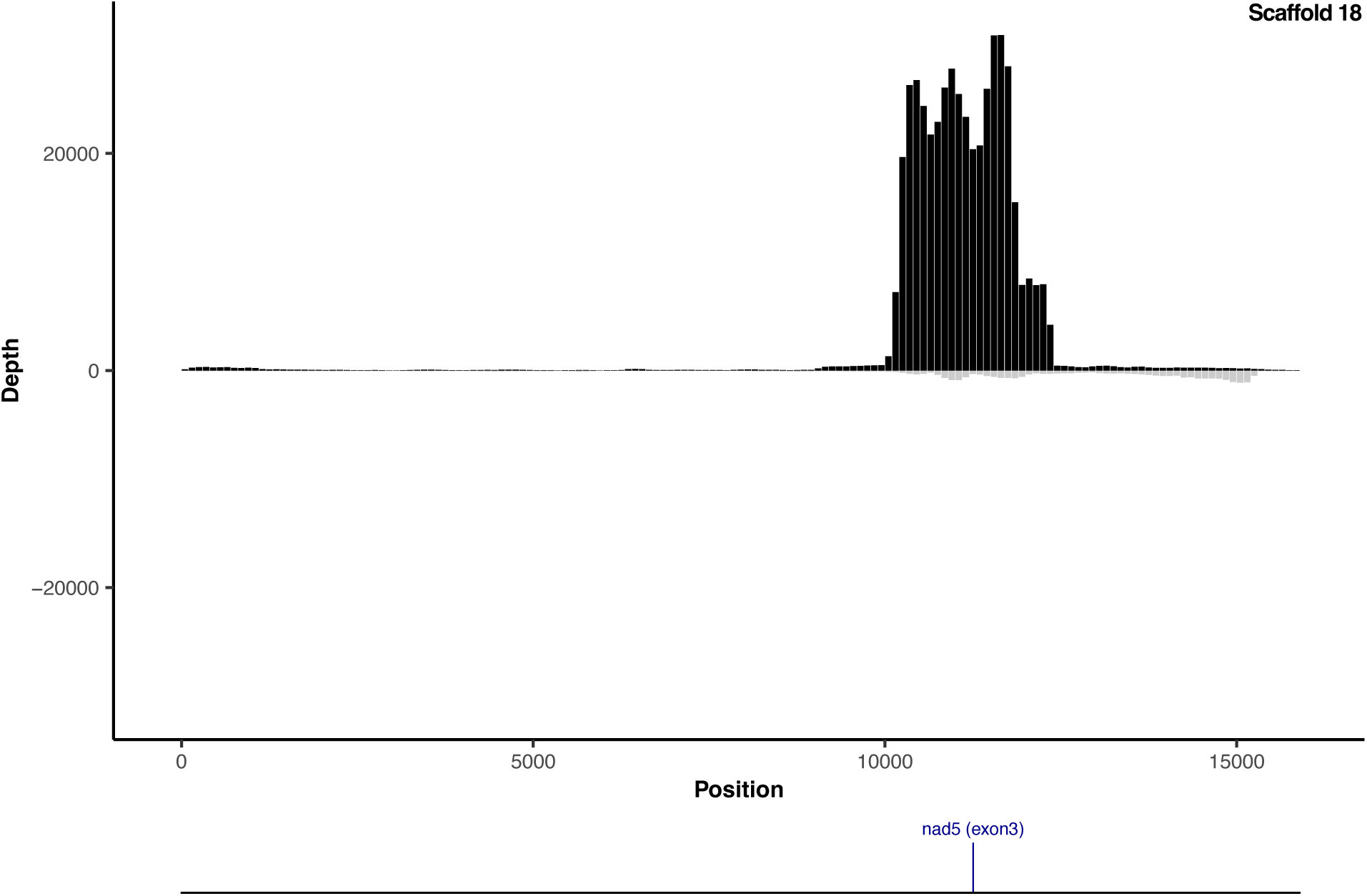
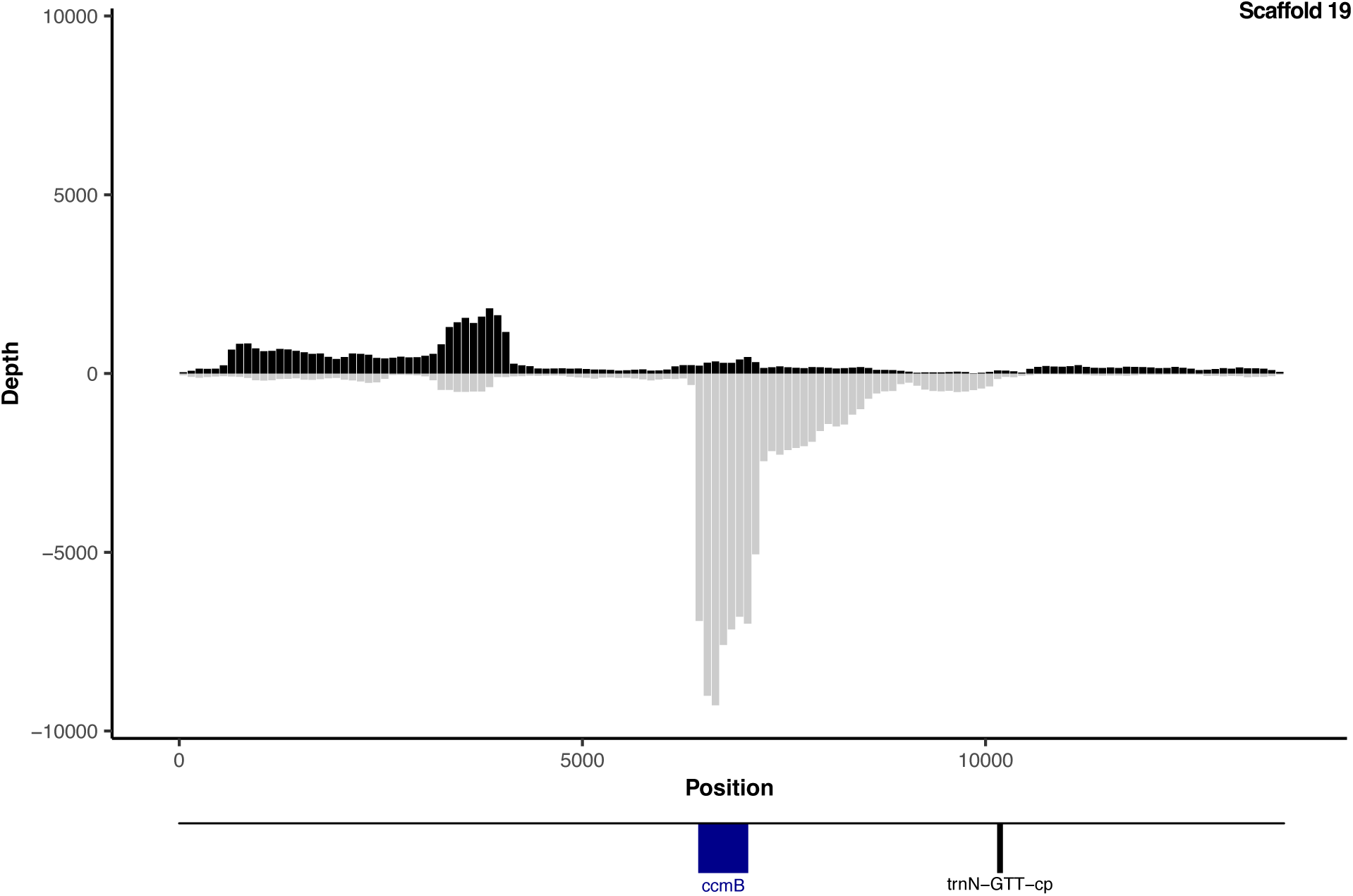
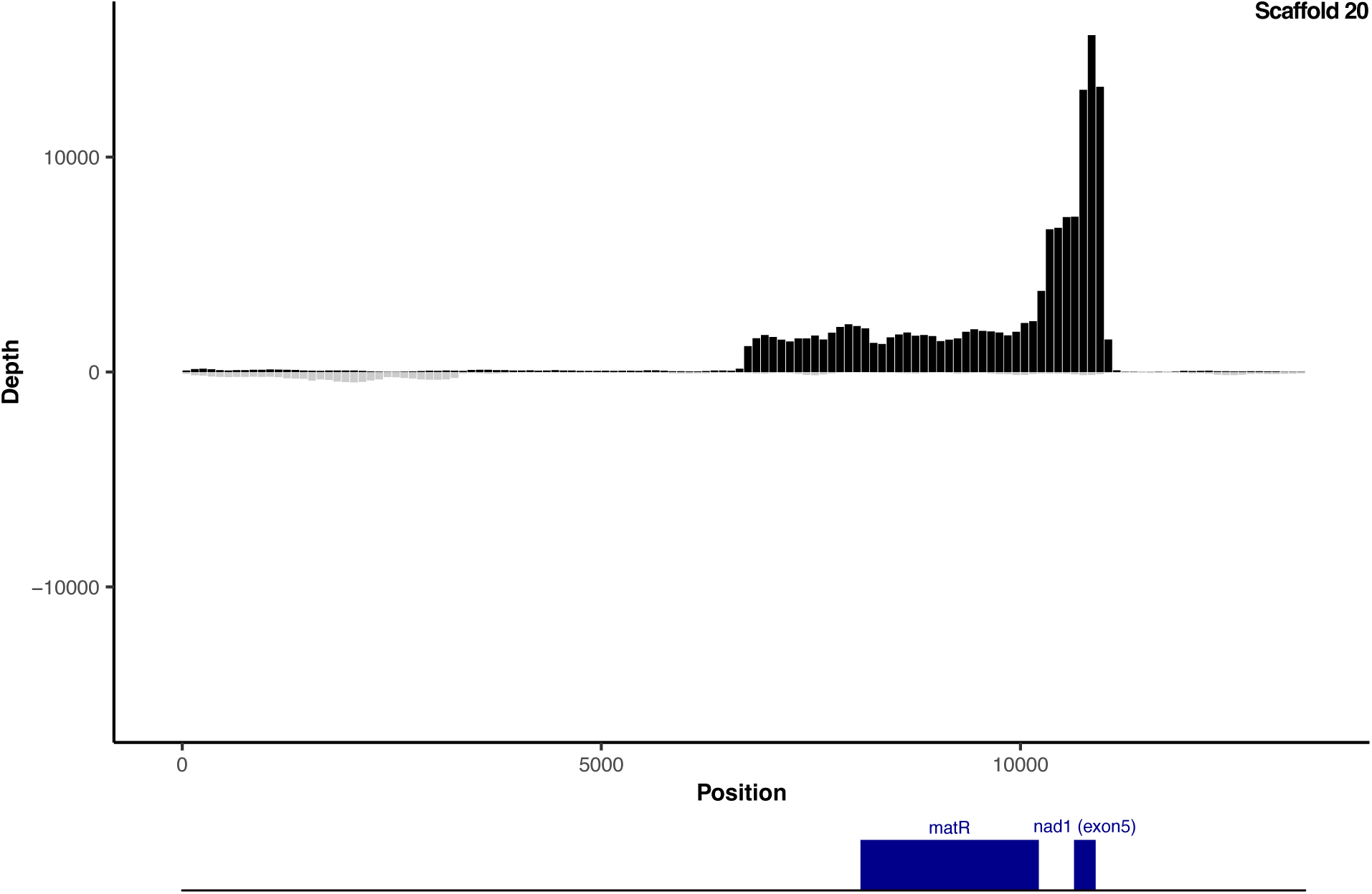
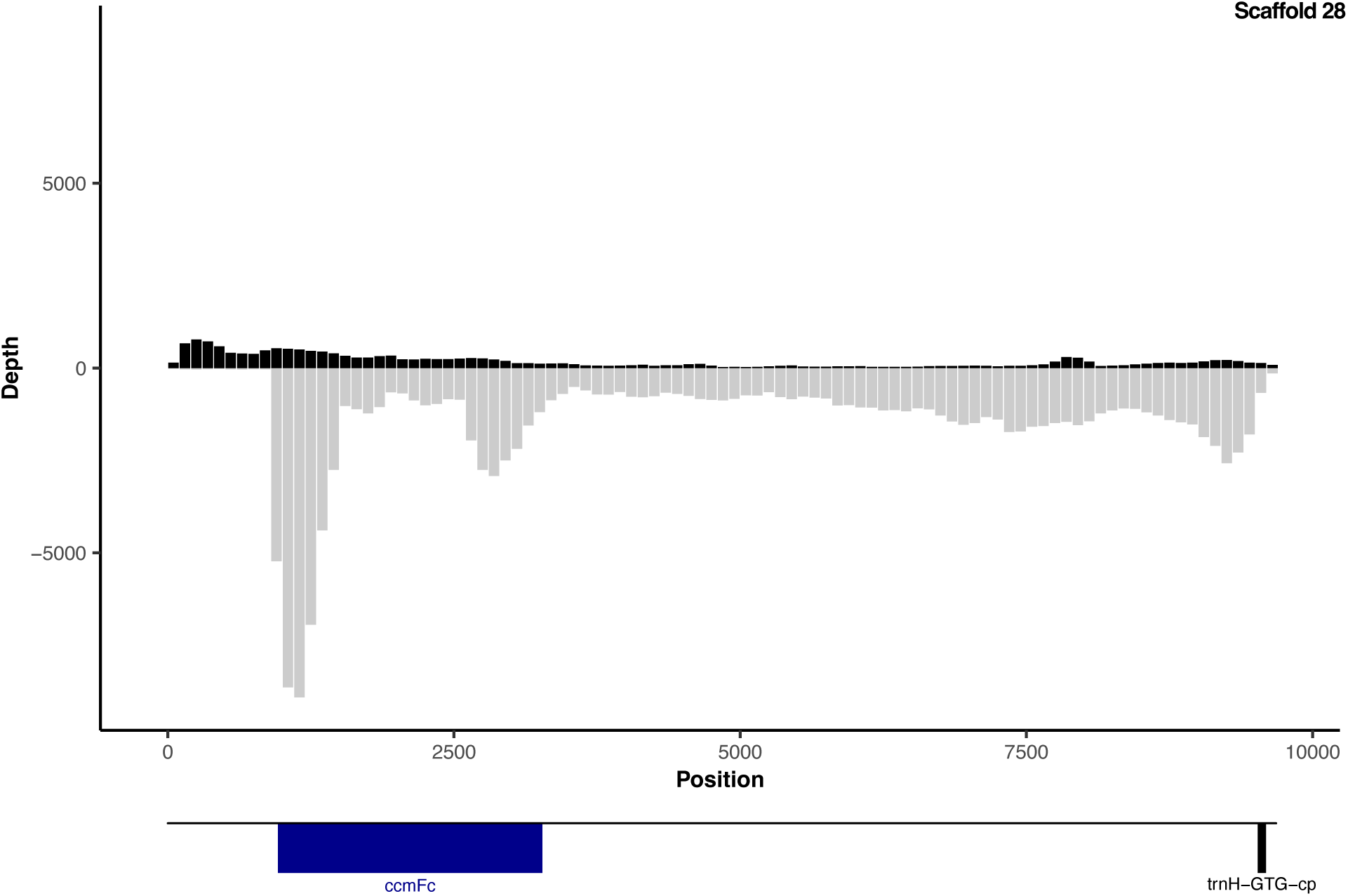
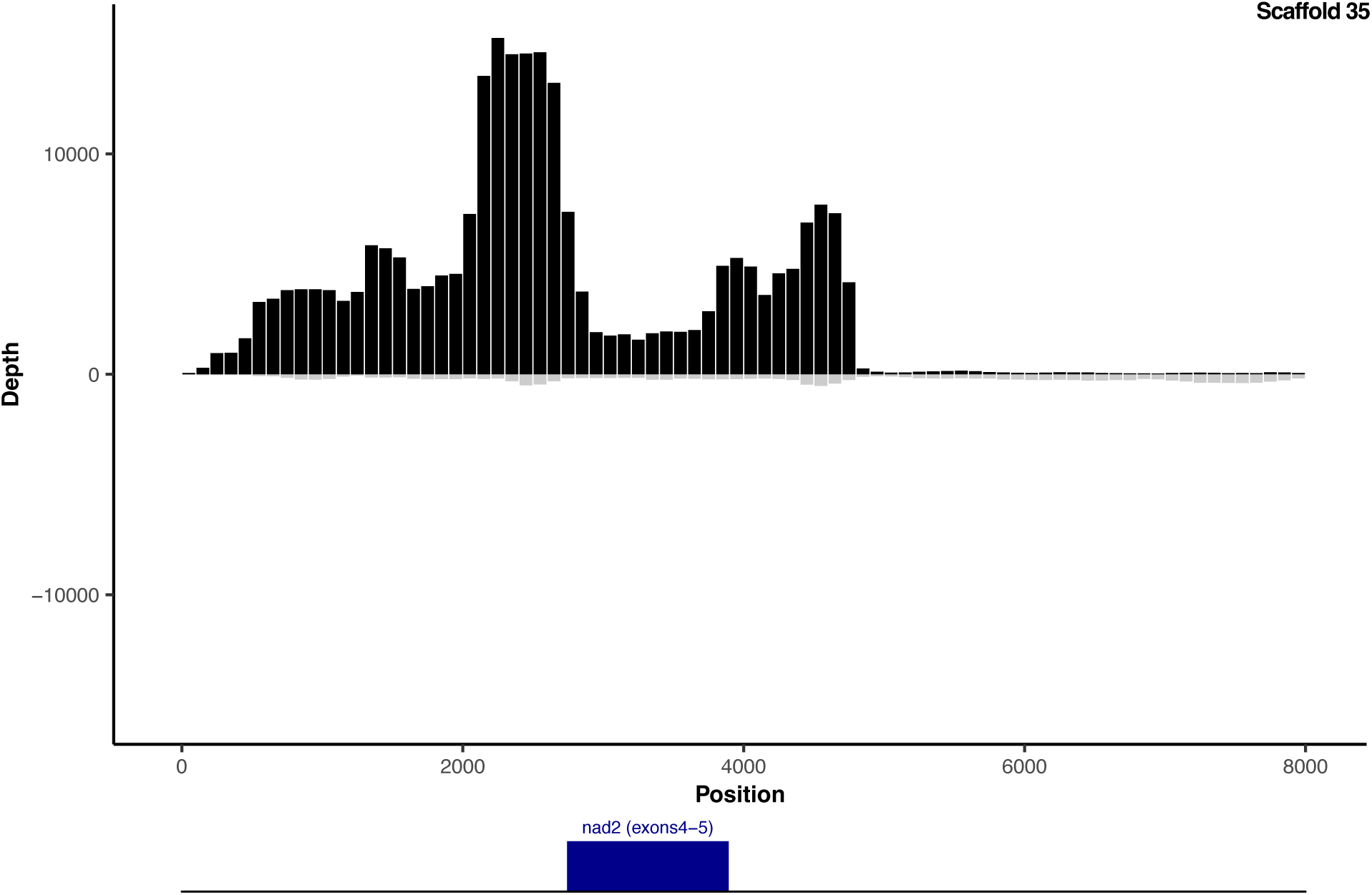
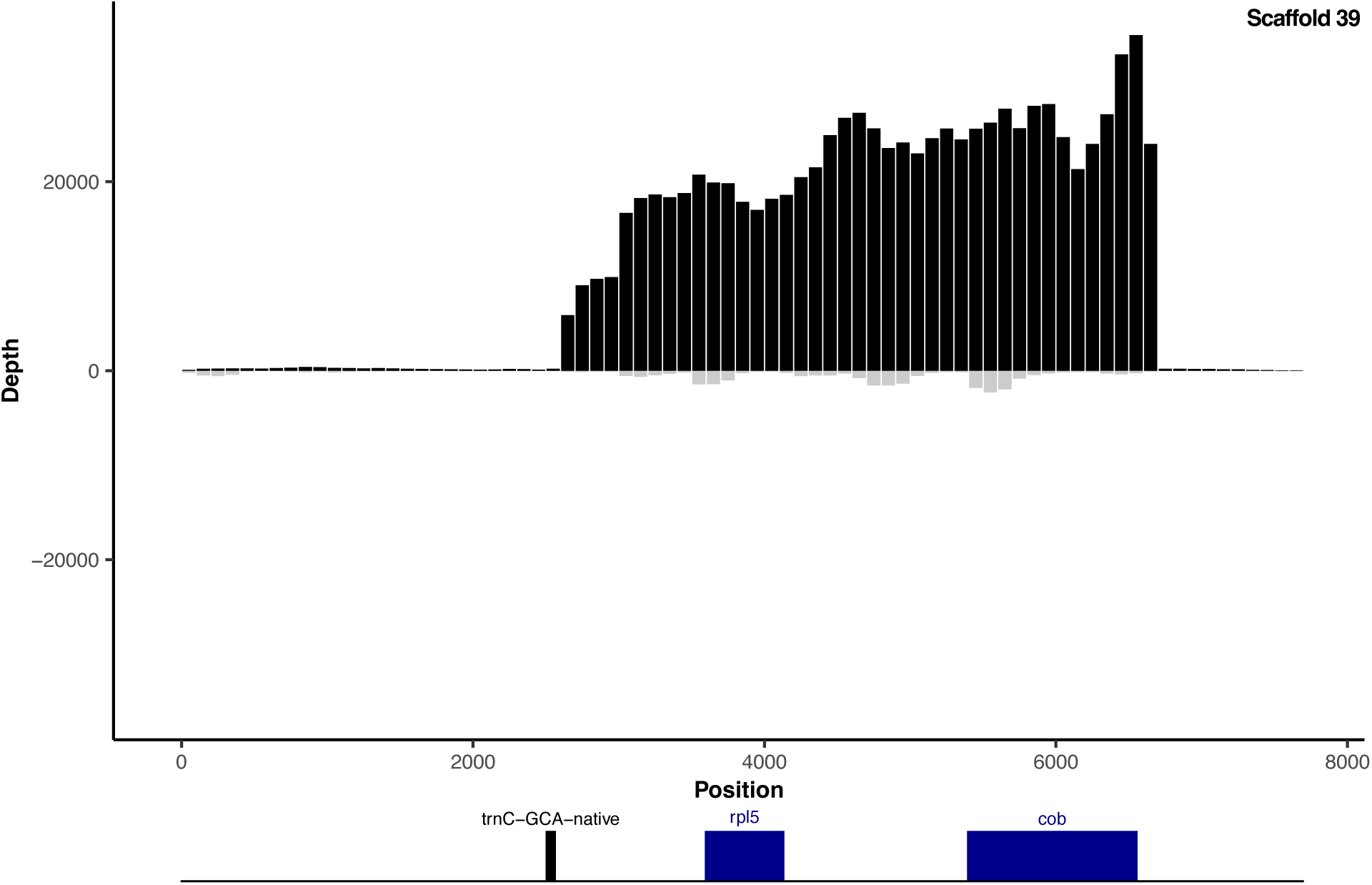
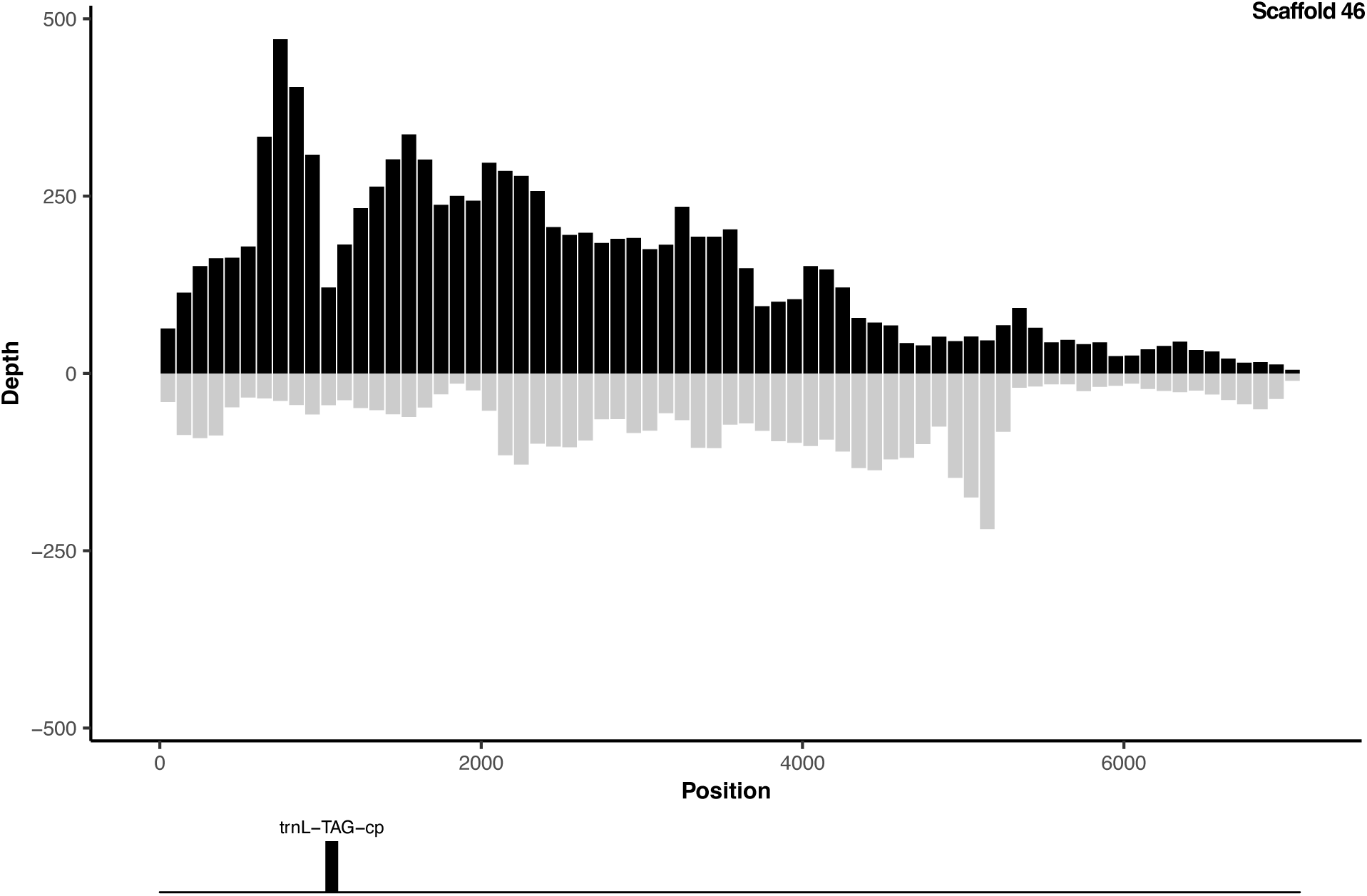
Transcript abundance measured from rRNA-depleted (Ribo-Zero) RNA-seq reads mapped to identified mitochondrial scaffolds from an assembly of total cellular DNA from *Corallorhiza maculata*. RNA and DNA were collected from the same individual (Plant 1). Depth is measured as average read sequence coverage (read count per position) for 100-bp windows across the length of the scaffolds. Positive and negative values indicate transcripts expressed in forward and reverse orientations relative to the reference sequence, respectively. Annotated genes are indicated in the diagram below the x-axis. Genes above and below the line correspond to being encoded on the forward and reverse strand of the reference sequence, respectively. The origins of tRNA gene are indicated as “native” (mitochondrial), “cp” (plastid intracellular gene transfer), and “bact” (bacterial horizontal gene transfer). Because of column-based RNA purification, library size selection, and the challenges inherent in sequencing tRNAs, these RNA-seq libraries are not expected to capture mature tRNA expression.

## References

Araiso Y, Huot JL, Sekiguchi T, Frechin M, Fischer F, Enkler L, Senger B, Ishitani R, Becker HD, Nureki O. 2014. Crystal structure of Saccharomyces cerevisiae mitochondrial GatFAB reveals a novel subunit assembly in tRNA-dependent amidotransferases. Nucleic Acids Res. 42:6052–6063.

Barrett CF, Freudenstein JV, Li J, Mayfield-Jones DR, Perez L, Pires JC, Santos C. 2014. Investigating the path of plastid genome degradation in an early-transitional clade of heterotrophic orchids, and implications for heterotrophic angiosperms. Mol. Biol. Evol. 31:3095–3112.

Behrens A, Rodschinka G, Nedialkova DD. 2021. High-resolution quantitative profiling of tRNA abundance and modification status in eukaryotes by mim-tRNAseq. Mol. Cell 81:1802–1815.e7.

von Braun SS, Sabetti A, Hanic-Joyce PJ, Gu J, Schleiff E, Joyce PBM. 2007. Dual targeting of the tRNA nucleotidyltransferase in plants: not just the signal. J. Exp. Bot. 58:4083–4093.

Camacho C, Coulouris G, Avagyan V, Ma N, Papadopoulos J, Bealer K, Madden TL. 2009. BLAST+: architecture and applications. BMC Bioinformatics 10:421.

Ceriotti LF, Warren JM, Virginia Sanchez-Puerta M, Sloan DB. 2024. The landscape of *Arabidopsis* tRNA aminoacylation. Plant J. In Press.

Chan PP, Lowe TM. 2019. TRNAscan-SE: Searching for tRNA genes in genomic sequences. Methods Mol. Biol. 1962:1–14.

Clark WC, Evans ME, Dominissini D, Zheng G, Pan T. 2016. tRNA base methylation identification and quantification via high-throughput sequencing. RNA 22:1771–1784.

Cognat V, Pawlak G, Pflieger D, Drouard L. 2022. PlantRNA 2.0: an updated database dedicated to tRNAs of photosynthetic eukaryotes. Plant J. 112:1112–1119.

Davidsen K, Sullivan LB. 2024. A robust method for measuring aminoacylation through tRNA-Seq. eLife 12:91554.2.

Dierckxsens N, Mardulyn P, Smits G. 2017. NOVOPlasty: de novo assembly of organelle genomes from whole genome data. Nucleic Acids Res. 45:e18.

Duchêne A-M, Giritch A, Hoffmann B, Cognat V, Lancelin D, Peeters NM, Zaepfel M, Maréchal-Drouard L, Small ID. 2005. Dual targeting is the rule for organellar aminoacyl-tRNA synthetases in *Arabidopsis thaliana*. Proceedings of the National Academy of Sciences 102:16484–16489.

Duchêne AM, Maréchal-Drouard L. 2001. The chloroplast-derived trnW and trnM-e genes are not expressed in Arabidopsis mitochondria. Biochem. Biophys. Res. Commun. 285:1213–1216.

Emanuelsson O, Nielsen H, Brunak S, von Heijne G. 2000. Predicting subcellular localization of proteins based on their N-terminal amino acid sequence. J. Mol. Biol. 300:1005–1016.

Evans ME, Clark WC, Zheng G, Pan T. 2017. Determination of tRNA aminoacylation levels by high-throughput sequencing. Nucleic Acids Res. 45:e133–e133.

Forner J, Weber B, Thuss S, Wildum S, Binder S. 2007. Mapping of mitochondrial mRNA termini in Arabidopsis thaliana: t-elements contribute to 5′ and 3′ end formation. Nucleic Acids Res. 35:3676–3692.

Frechin M, Senger B, Brayé M, Kern D, Martin RP, Becker HD. 2009. Yeast mitochondrial Gln-tRNA(Gln) is generated by a GatFAB-mediated transamidation pathway involving Arc1p-controlled subcellular sorting of cytosolic GluRS. Genes Dev. 23:1119–1130.

Grabherr MG, Haas BJ, Yassour M, Levin JZ, Thompson DA, Amit I, Adiconis X, Fan L, Raychowdhury R, Zeng Q, et al. 2011. Full-length transcriptome assembly from RNA-Seq data without a reference genome. Nat. Biotechnol. 29:644–652.

Joyce PB, Gray MW. 1989. Chloroplast-like transfer RNA genes expressed in wheat mitochondria. Nucleic Acids Res. 17:5461–5476.

Kitazaki K, Kubo T, Kagami H, Matsumoto T, Fujita A, Matsuhira H, Matsunaga M, Mikami T. 2011. A horizontally transferred tRNACys gene in the sugar beet mitochondrial genome: evidence that the gene is present in diverse angiosperms and its transcript is aminoacylated. Plant J. 68:262–272.

Knie N, Polsakiewicz M, Knoop V. 2015. Horizontal gene transfer of chlamydial-like tRNA genes into early vascular plant mitochondria. Mol. Biol. Evol. 32:629–634.

Langmead B, Salzberg SL. 2012. Fast gapped-read alignment with Bowtie 2. Nat. Methods 9:357–359.

Li H, Handsaker B, Wysoker A, Fennell T, Ruan J, Homer N, Marth G, Abecasis G, Durbin R. 2009. The Sequence Alignment/Map format and SAMtools. Bioinformatics 25:2078–2079.

Marchfelder A, Schuster W, Brennicke A. 1990. In vitro processing of mitochondrial and plastid derived tRNA precursors in a plant mitochondrial extract. Nucleic Acids Res. 18:1401–1406.

Martin M. 2011. Cutadapt removes adapter sequences from high-throughput sequencing reads. EMBnet.journal 17:10–12.

Nurk S, Meleshko D, Korobeynikov A, Pevzner PA. 2017. metaSPAdes: a new versatile metagenomic assembler. Genome Res. 27:824–834.

Padhiar NH, Katneni U, Komar AA, Motorin Y, Kimchi-Sarfaty C. 2024. Advances in methods for tRNA sequencing and quantification. Trends Genet. 40:276–290.

Pujol C, Bailly M, Kern D, Marechal-Drouard L, Becker H, Duchene AM. 2008. Dual-targeted tRNA-dependent amidotransferase ensures both mitochondrial and chloroplastic Gln-tRNAGln synthesis in plants. Proceedings of the National Academy of Sciences 105:6481–6485.

Ramírez SR, Gravendeel B, Singer RB, Marshall CR, Pierce NE. 2007. Dating the origin of the Orchidaceae from a fossil orchid with its pollinator. Nature 448:1042–1045.

Rice DW, Alverson AJ, Richardson AO, Young GJ, Sanchez-Puerta MV, Munzinger J, Barry K, Boore JL, Zhang Y, dePamphilis CW, et al. 2013. Horizontal transfer of entire genomes via mitochondrial fusion in the angiosperm Amborella. Science 342:1468–1473.

Richardson AO, Rice DW, Young GJ, Alverson AJ, Palmer JD. 2013. The “fossilized” mitochondrial genome of Liriodendron tulipifera: ancestral gene content and order, ancestral editing sites, and extraordinarily low mutation rate. BMC Biol. 11:29.

Sanchez-Puerta MV, García LE, Wohlfeiler J, Ceriotti LF. 2017. Unparalleled replacement of native mitochondrial genes by foreign homologs in a holoparasitic plant. New Phytol. 214:376–387.

Schneider A. 2011. Mitochondrial tRNA import and its consequences for mitochondrial translation. Annu. Rev. Biochem. 80:1033–1053.

Schweizer U, Bohleber S, Fradejas-Villar N. 2017. The modified base isopentenyladenosine and its derivatives in tRNA. RNA Biol. 14:1197–1208.

Sinn BT, Barrett CF. 2020. Ancient mitochondrial gene transfer between fungi and the orchids. Mol. Biol. Evol. 37:44–57.

Small I, Akashi K, Chapron A, Dietrich A, Duchene AM, Lancelin D, Maréchal-Drouard L, Menand B, Mireau H, Moudden Y. 1999. The strange evolutionary history of plant mitochondrial tRNAs and their aminoacyl-tRNA synthetases. J. Hered. 90:333–337.

Suzuki T. 2021. The expanding world of tRNA modifications and their disease relevance. Nat. Rev. Mol. Cell Biol. 22:375–392.

Valencia-D J, Neubig KM, Clark DP. 2023. The origin and fate of fungal mitochondrial horizontal gene transferred sequences in orchids (Orchidaceae). Bot. J. Linn. Soc. 203:162–179.

Warren JM, Salinas-Giegé T, Hummel G, Coots NL, Svendsen JM, Brown KC, Maréchal-Drouard L, Sloan DB. 2021. Combining tRNA sequencing methods to characterize plant tRNA expression and post-transcriptional modification. RNA Biol. 18:64–78.

Warren JM, Salinas-Giegé T, Triant DA, Taylor DR, Drouard L, Sloan DB. 2021. Rapid shifts in mitochondrial tRNA import in a plant lineage with extensive mitochondrial tRNA gene loss. Mol. Biol. Evol. 38:5735–5751.

Warren JM, Sloan DB. 2020. Interchangeable parts: The evolutionarily dynamic tRNA population in plant mitochondria. Mitochondrion 52:144–156.

Watkins CP, Zhang W, Wylder AC, Katanski CD, Pan T. 2022. A multiplex platform for small RNA sequencing elucidates multifaceted tRNA stress response and translational regulation. Nat. Commun. 13:2491.

Wicke S, Naumann J. 2018. Molecular evolution of plastid genomes in parasitic flowering plants. In: Advances in Botanical Research. Vol. 85. Elsevier. p. 315–347.

Wilusz JE. 2015. Removing roadblocks to deep sequencing of modified RNAs. Nat. Methods 12:821–822.

